# Long-read sequencing characterizes mitochondrial and plastid genome variants in Arabidopsis *msh1* mutants

**DOI:** 10.1101/2022.02.28.481893

**Authors:** Yi Zou, Weidong Zhu, Daniel B. Sloan, Zhiqiang Wu

## Abstract

The abundant repeats in plant mitochondrial genomes can cause rapid genome rearrangements and are also a major obstacle in short-read sequencing studies. Nuclear-encoded proteins such as MSH1 are known to suppress the generation of repeat-associated mitochondrial genome variants, but our understanding of these mechanisms has been constrained by the limitations of short-read technologies. Here, we used highly accurate long-read sequencing (PacBio HiFi) to characterize mitochondrial and plastid genome variants in *Arabidopsis thaliana msh1* mutant individuals. The HiFi reads provided a global view of recombination dynamics with detailed quantification of parental and crossover recombination products for both large and small repeats. We found that recombination breakpoints were distributed relatively evenly across the length of repeated sequences and detected widespread internal exchanges of sequence variants between pairs of imperfect repeats in the mitochondrial genome of *msh1* mutants. Long-read assemblies of mitochondrial genomes from seven other *Arabidopsis thaliana* wild-type accessions differed by repeat-mediated structural rearrangements similar to those observed in *msh1* mutants, but they were all in a simple low-heteroplasmy state. The *Arabidopsis* plastid genome generally lacks small repeats and exhibited a very different pattern of variant accumulation in *msh1* mutants compared with the mitochondrial genome. Our data illustrate the power of HiFi technology in studying repeat-mediated recombination in plant organellar genomes and improved the sequence resolution for recombinational processes suppressed by MSH1.

**Significance:** Plant organellar genomes can undergo rapid rearrangements. Long-read sequencing provides a detailed and quantitative view of mitochondrial and plastid genome variants normally suppressed by MSH1, advancing our understanding of plant organellar genome dynamics.

## Introduction

Mitochondria and plastids are major energy converters as well as central metabolic compartments of plant cells. In plants, mitochondrial (mt) and plastid (pt) genomes have followed very different evolutionary trajectories. Compared with the rather conserved structures of pt genomes (Lee et al., 2016), plant mt genomes exhibit dramatic variation among species (Wu et al., 2022). Although coding sequences in angiosperm mt genome have very low substitution rates, their gene order and intergenic sequences are rarely conserved (Wolfe et al., 1987; Palmer and Herbon 1988; Christensen, 2021). Angiosperm mt genomes typically contain numerous non-tandem repeats, and recombination between different copies of repeat sequences causes a high level of structural variation (Wynn and Christensen, 2019). These repeats show little or no sequence conservation between species and are often grouped based on size ranges (Arrieta-Montiel et al., 2009). Large repeats (≥1000 bp) are usually identical in sequence and recombine at high frequency, leading to isomerization of the genome (Unseld et al., 1997). The recombination frequencies of intermediate-sized repeats (50-1000 bp) are lower but sensitive to genetic, environmental and developmental triggers that make them frequent enough to have important phenotypic effects (Gualberto and Newton 2017; Mackenzie and Kundariya, 2020). Shorter repeats (<50 bp) typically do not show recombination activity, but still play important roles during evolution. These short sequences are often described as “microhomologies”, but the very shortest of them (e.g., <10 bp) are unlikely to reflect true homology, because such similarities can easily occur at random. Despite their importance, repeats in sequenced mt genomes are not always annotated correctly or completely even in model plant species (Wynn and Christensen, 2019). Moreover, these repeats may produce complex assembly graphs, which hinder a clear characterization of mt genomes (Masutani et al., 2021). Thus, mechanisms underlining the repeats and their roles in destabilizing plant mt genomes require a more thorough exploration.

Recombination in plant mt genomes is likely involved in repairing DNA damage, such as double-strand breaks (DSBs), which can disrupt genome integrity and prevent the progression of replication and transcription activities. To deal with DSBs, two main branches of repair systems exist: homology-directed repair (HDR) and non-homologous end joining (NHEJ) (Hussmann et al., 2021). HDR requires homologous dsDNA templates for new strand synthesis. HDR involving both DSB ends will lead to non-crossover repair products, which is a process known as gene conversion. In contrast, HDR involving only one DSB end and non-allelic templates is known as break-induced repair and leads to crossover products (Christensen, 2013).

Nuclear genes encode a set of mt- and mt-targeted DNA binding proteins that are involved in DNA recombination, replication, and repair (RRR; Gualberto and Newton, 2017; Chevignv et al., 2020; Fuchs et al., 2020). Some RRR genes, such as those in the *RECA*, *POLI*, and *SSB* families, have conserved roles, while others are evolutionary innovations. Unlike bacterial MutS counterparts, the dual-targeted (mt and pt) MSH1 enzyme in plants combines a mismatch-recognition domain (MutS I) and a predicted DSB-generating GIY-YIG endonuclease domain in a single protein (Abdelnoor et al., 2006). MSH1 has been hypothesized to participate in novel DNA repair mechanisms in plant organelles (Christensen, 2013; Ayala-Garcia et al., 2018; Fukui et al, 2018; Wu et al., 2020; Wynn et al., 2020).

Arabidopsis *msh1* (=*chm*) mutants were originally isolated based on a green-white leaf variegation phenotype (Rédei, 1973). Early studies focused on reproducible mt genome rearrangements at a specific region near the *atp9* gene in *msh1* mutants and explored processes such as asymmetric recombination and substoichiometric shifting (Martínez-Zapater et al., 1992; Sakamoto et al., 1996; Abdelnoor et al., 2003; Shedge et al., 2007). They showed that new bands in DNA gel blots in *msh1* mutants were similar to bands observed in other Arabidopsis accessions, which existed at substoichiometric levels in wild-type individuals and could be detected by PCR. Detecting pt genome rearrangements in *msh1* mutants by DNA gel blots was more difficult and only successful in dissected white tissues using probes to detect rearrangements already known in another mutant (Xu et al., 2011). The more limited structural effects in the Arabidopsis pt genome may reflect a relative lack of dispersed repeats. An analysis of *msh1* mutants in the moss *Physcomitrella patens* has revealed more extensive pt genome variation associated with repeat-mediated recombination (Odahara et al., 2017).

The first systematic analysis across the mt genome in *msh1* mutants confirmed asymmetric recombination at 23 groups of repeats using probes corresponding to 33 near-perfect intermediate-sized repeats which were inferred from blast analysis of the Arabidopsis C24 mt genome (Arrieta-Montiel et al., 2009). The intensity of bands detected by a specific probe varied among *msh1* individuals and generations. In some cases, weak bands corresponding to secondary recombination (multiple rearrangements) and loss of parental bands were observed (Shedge et al., 2007; Arrieta-Montiel et al., 2009). It was also reported that lower *MSH1* expression caused mt rearrangements that produced cytoplasmic male sterile (CMS) lines in tomato and allowed fertility reversion of lines of *Brassica juncea* (Sandhu et al., 2007; Zhao et al., 2016, 2021). These studies demonstrated that MSH1 maintains the structural integrity of the mt genome through suppressing illegitimate recombination at repeats. We also recently showed increased frequencies of de novo point mutations in organelle genomes of *msh1* mutants, suggesting a role of MSH1 in maintaining lower mutation rates in organellar genomes compared with the nuclear genome (Wu et al., 2020).

Despite this extensive work on *MSH1* in Arabidopsis, we still lack a complete understanding of its effect on repeat dynamics, in part due to the technical difficulties of traditional DNA gel blots, quantitative PCR, and short-read sequencing. For example, a previous study classified mt genome rearrangements in Arabidopsis *msh1* mutants using early-generation Illumina sequencing (Davila et al. 2011). This work represented a key advance in the genome-wide characterization of mt recombinational activity, including sequence-level resolution of some recombinant molecules. However, the short read-lengths (36-bp) available at the time cannot span entire repeats including the regions of perfect sequence identity that often separate internal variants in imperfect repeats, placing significant limitations on analysis of recombination breakpoints. Short-read sequencing also precludes detection of more complex structural rearrangements resulting from recombination at multiple different repeats. Davila et al. (2011) did not detect recombinationally active repeats smaller than 50 bp. They proposed that structural differences among Arabidopsis accessions were mainly the result of repeat-mediated recombination, microhomology-mediated end joining (MMEJ), and NHEJ, but they did not detect MMEJ or NHEJ activities with short reads in *msh1* mutants. Davila et al. (2011) also documented discontinuous patterns of single-nucleotide variants (SNVs) and indels that were exchanged between pairs of imperfect repeats. Once again, however, the resolution of this important analysis was limited due to short read lengths, and it was only applied to a subset of repeat pairs. Fortunately, the advent of long-read sequencing technologies has created the opportunity to analyze structural variants in plant mt genome assemblies with much more accuracy and scalability (Kozik et al., 2019; Yu et al., 2022).

In this work, we aim to comprehensively investigate mt and pt genome instabilities in Arabidopsis *msh1* mutants. Using PacBio high-fidelity (HiFi) data for individual plants, we identified structural variants resulting from one or more repeat-mediated rearrangements, as well as lower frequency MMEJ and NHEJ events. We provide a global map of mt genome instabilities with a more detailed and accurate characterization of different repeats. For imperfect repeats, we systematically analyzed the distribution of breakpoints and observed the exchange of sequence variants in molecules that did not exhibit any structural rearrangements, which was an unexpected process suppressed by MSH1. We also show that mt genomes of seven *Arabidopsis thaliana* accessions are all in a low-heteroplasmy states, despite differing by the same types of rearrangements observed in *msh1* mutants. The total structural variation was very low in pt genomes. But some variants show individual-specific accumulation in *msh1* mutants, suggesting a strong influence of random heteroplasmic sorting. These results are important for improving mt and pt genome-editing technologies and understanding the distinct patterns of evolution in these organellar genomes.

## Results

### Mapping of HiFi reads identified structural variants in Arabidopsis mitochondrial genomes

We generated F_3_ families from a cross between wild type Arabidopsis Columbia-0 (Col-0) and an *msh1* mutant line (*chm1-1*, CS3372) in an earlier project (Wu et al., 2020). To analyze structural rearrangements in the organellar genomes in homozygous *msh1* mutants, we collected aboveground tissue from 11 individual plants (10 *msh1* individuals: M3-17-1, M3-17-2, M3-17-4, M3-17-7, M3-17-8, M3-17-9, M3-43-2, M3-43-4, M3-43-5, M3-43-6; 1 wild-type individual: W3-5-2) from 3 different F_3_ families (Table S1; Figure S1) and prepared separate libraries for each plant for PacBio HiFi sequencing. The libraries generated a total of 34.36 Gb of HiFi reads (12.7-18.1 kb mean read length). These reads were first mapped to the Col-0 reference genome by minimap2 (Li, 2018) to separate nuclear, pt and mt proportions. After excluding one outlier with very low yield (M3-17-2), the coverage of the nuclear genome ranged from 8.5× to 46.4×, the coverage of the pt genome ranged from 899.0× to 3093.8×, and the coverage of the mt genome ranged from 30.9× to 247.9× (Table S2). The pt (4.96%-12.85%) and mt (0.93%-2.13%) proportions were used for downstream analysis (Figure S2; Table S3 and S4).

Published mt genomes are usually represented as a linearized sequence of the “master circle”. However, various sub-stoichiometric isoforms exist due to frequent repeat-mediated structural rearrangements (Kozik et al., 2019). Structural rearrangements are indicated by reads whose parts map to discontinuous positions in the reference genome (Figure 1a). We developed a pipeline to detect these structural rearrangements (available at https://github.com/zouyinstein/hifisr), in which mt genome-mapped reads were aligned against the reference sequence (NC_037304.1; Sloan et al., 2018) via the NCBI blastn software. Using custom Python scripts, we grouped the reads by discontinuity and mapping coordinates, and calculated alignment overlap (AO) length (i.e., repeat length) and read counts for each group (Table S5). After additional manual checking and curation, we were able to distinguish reads with no rearrangements (zero-rearrangement reads) from those containing one, two, three, or four repeat-mediated rearrangements (≥50 bp) in a single read (Figure 1b; Table S6). In wild type, 7.50% of total reads accounted for rearrangements mediated by non-tandem repeats, most of which were associated with the two large repeat pairs (Figure 1c). In *msh1* mutants, this ratio increased to 31.63% of total reads, containing increased proportions of reads with rearrangements mediated by intermediate-sized repeats and reads with multiple rearrangements (Figure 1b,c; Table S7 and S8). A few low-frequency events were also detected, including MMEJ (AO length between 2 and 49 bp) and NHEJ (AO length of 0 or 1 bp), replication slippage associated with tandem repeats, and insertions (poly-G/C tracts in most cases) (Figure 1b; Table S6). In particular, the proportion of MMEJ events also increased in *msh1* mutants compared with wild-type (Figure 1b). Taken together, these data indicate that mt genome structural variants in *msh1* mutants is mainly (> 96.7%) associated with non-tandem repeats. Compared to short-read sequencing, our results open the opportunity for investigating multiple rearrangements in a single read and low-frequency categories of structural variants.

**Figure 1.**
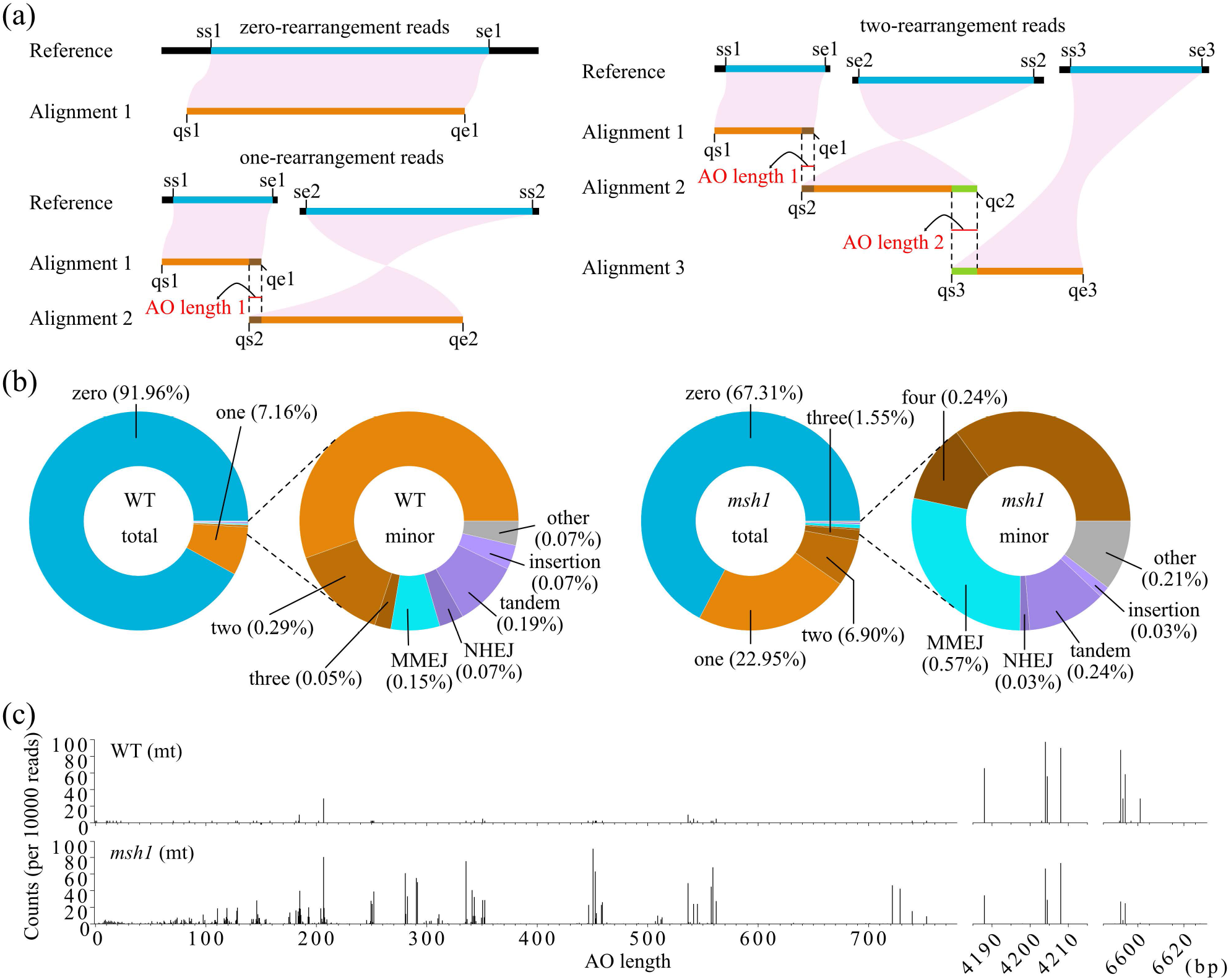
Mapping of HiFi reads identified structural variants in Arabidopsis mt genomes (a) Schematics show read alignment to the reference genome by blastn. Reads mapped in their entirety to one position indicate zero-rearrangement events. Reads mapped to two discontinuous positions indicate a single rearrangement. Start and end coordinates of blastn queries are shown as qs1, qe1 for Alignment 1, and qs2, qe2 for Alignment 2. Start and end coordinates of corresponding blastn subjects are shown as ss1, se1 for Alignment 1, and ss2, se2 for Alignment 2. Alignment overlap length (AO length) 1 in red color equals “qe1 – qs2 + 1”. The overlapping regions of two alignments of succussive parts of the same read correspond to sequence homologies such as non-tandem repeats. Coordinates of se1 and ss2 in the genome correspond to repeat boundaries of the rearrangement event. Reads mapped to three discontinuous positions indicate two rearrangements in a single read. (b) Plots are proportions of different types of mt genome rearrangements detected by HiFi reads in wild-type and *msh1* mutants; zero, one, two, three, and four denote ratios of zero-, one-, two-, three-, four-rearrangement events; MMEJ, microhomology-mediated end joining (2-49 bp homology sequence size); NHEJ, non-homologous end-joining (0-1 bp homology sequence size); tandem, large-scale tandem repeats-mediated variants; insertion, large insertions (mostly poly-G and poly-C tracts). The percentages of rearrangement categories are shown in parentheses. The total circles represent all reads, and the minor circles represent the last 2% of total reads to show low-frequency categories. (c) The distribution of AO length in mt one-rearrangement events detected by HiFi reads. Counts from read groups with equal AO lengths are summed. Reads from different *msh1* individuals are merged. Read counts are normalized to 10000 total mt genome-mapped reads.

### Global patterns of mt genome variants in *msh1* mutants

By visualizing all the junctions and read counts of one-rearrangement reads, we found that rearrangements in wild-type individuals were mainly associated with the Large1, Large2, Q, and V repeats, as well as a few other repeats at very low frequency (Figure S3). In *msh1* mutants, the activity of increased mt genome rearrangements was not randomly distributed (Figure 2a). Instead, they were mostly associated with the subset of repeats that were named and annotated in previous studies with the most extreme variability observed at three regions containing large repeats and two additional regions with intermediate-sized repeats (Figure S4; Arrieta-Montiel et al., 2009; Davila et al., 2011; Sloan et al., 2018). With more exhaustive blastn parameters, we found that many smaller repeats that were not previously named were also associated with mt genome rearrangements (smaller unlabeled dots in Figure 2a and S3; Table S9). However, rearrangements mediated by these small repeats were supported by very few reads and were detected in fewer samples (Figure 2b,c).

**Figure 2.**
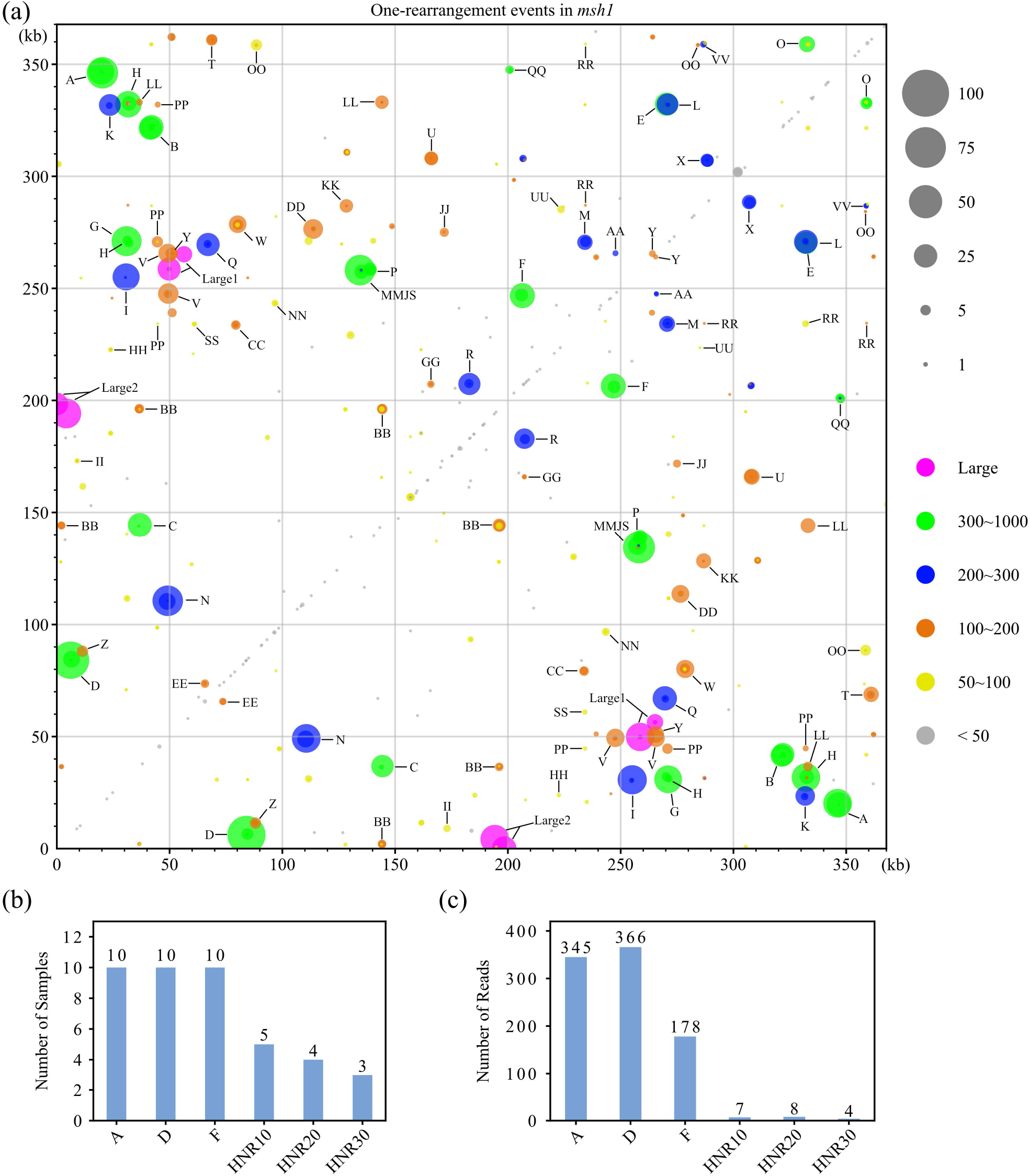
Patterns of one-rearrangement mt reads detected in *msh1* mutants (a) Dot plots show global patterns of the junctions and relative read counts of mt one-rearrangement reads in *msh1* mutants. Signals from previously named repeats are labeled. In *msh1*, recombination events mediated by intermediate-sized repeats are common. Reads from different *msh1* individuals are merged. There are 5544 one-rearrangement reads out of 24162 total reads in all *msh1* samples. Colors represent categories of alignment overlap length (AO length). The x position of each dot is the end coordinate of the blastn subject for Alignment 1 (se1). The y position of each dot is the start coordinate of the blastn subject for Alignment 2 (ss2). The sizes of dots denote read counts normalized to 10000 total mt genome-mapped reads per sample. The near symmetrical appearance of the plot is a result of HiFi reads aligned in both forward and reverse directions relative to the reference genome. The spots in mirrored positions should not be interpreted as quantifications of the two reciprocal recombination products. (b, c) Rearrangements mediated by newly named repeats are supported by (c) fewer reads and are observed in (b) fewer samples.

Unlike previous DNA gel blots or PCR-based analysis, we could compare the activities of different repeats. According to relative recombination frequencies, 18 groups (Large1, Large2, MMJS, A, B, C, D, E, F, G, N, I, L, Q, R, H, V, BB) showed high activity (read count ≥ 50 in *msh1*, normalized to 10000 total mt genome-mapped reads), 17 groups (P, O, QQ, M, K, X, U, T, W, CC, Z, DD, Y, LL, HNR37, KK, OO) showed intermediate activity (read count of 10 to 49 in *msh1*) and the 64 remaining groups showed low activity (Figure S5a; Table S10-S13). The factors that contributed to different recombination frequencies were not limited to repeat length, identity or copy number, because some repeats still fell far from the regression line when we considered all two-copy repeats with high sequence identity (mismatches + gaps ≤ 2; Figure S5b and Table S14) and different pairs within the same three-copy repeat group showed different recombination frequencies (Table S15).

Asymmetrical recombination in *msh1* mutants was first reported based on DNA gel blot analysis and was also estimated based on read depth across one of the repeat boundaries (Arrieta-Montiel et al., 2009; Davila et al., 2011). Using long reads across both repeat boundaries for a given repeat group, we could estimate asymmetrical recombination events in more detail (Figure S6). The polarity of asymmetrical recombination between the two *msh1* mutant families was largely conserved, with small differences in intensity and accompanied loss of parental forms for some repeat groups which could be caused by heteroplasmic sorting (Figure 3; Table S16). Notably, we observed more pronounced asymmetrical recombination and severe loss of one parental form (2-2) for Large1 in *msh1* mutants compared with wild-type (Figure 3). Such large repeats are typically found to recombine symmetrically in the literature (Figure 3; Arrieta-Montiel et al., 2009).

**Figure 3.**
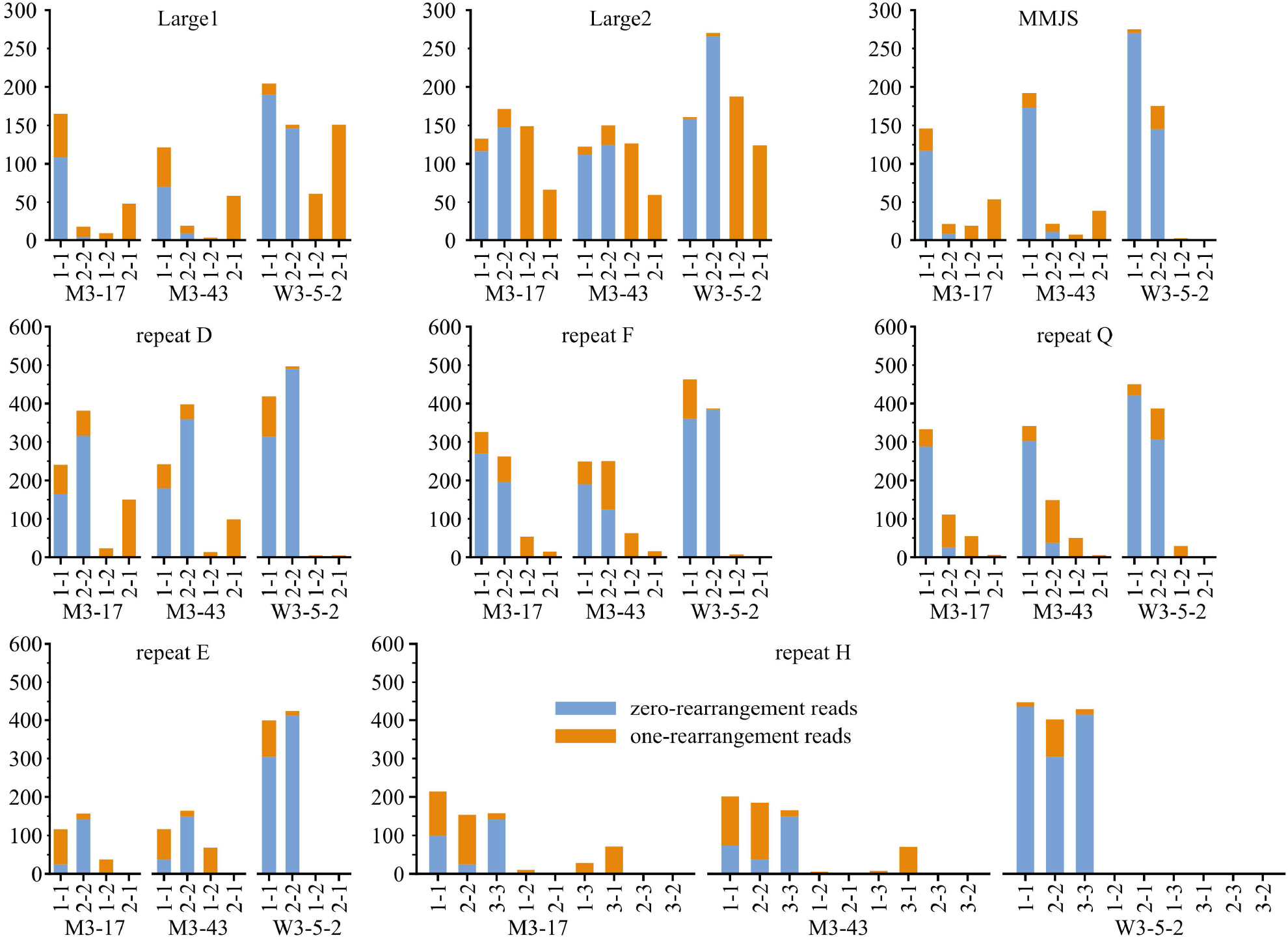
Asymmetrical recombination events and accompanied loss of parental forms Stacked bar plots show HiFi read counts indicating parental forms (1-1, 2-2, 3-3) and crossover recombination products (1-2, 2-1, 1-3, 3-1, 2-3, 3-2) of representative repeats (Large1, Large2, MMJS, D, F, Q, E, H) in wild-type and *msh1* mutants. The unequal counts 1-2 versus 2-1 or 1-3 versus 3-1 in each repeat group indicate asymmetrical recombination or simply preferential accumulation of only one of the crossover products, which is accompanied by a striking loss of 2-2 versus 1-1 parental forms in Large1, MMJS, and Q. Due to the locations of H-2 and H-3 within E-1 and E-2, recombination events between H-2 and H-3 were assigned to the larger two-copy repeats E-1 and E-2 and, thus, appear as zero values in this plot. Both zero-rearrangement reads (blue bars) and one-rearrangement reads (orange bars) were used for calculation. Read counts are normalized to 10000 total mt genome-mapped reads per sample. Reads from different *msh1* families are merged.

Long reads enabled the identification of rearrangements that involve two or more groups of repeats. For example, the increased two-rearrangement reads involving Large1 and Large2 indicated more fragmented and heteroplasmic mt genomes in *msh1* mutants compared with wild-type (Figure S7). Together, our data provide a much more detailed and accurate view of mt genome instabilities of *msh1* mutants with a quantitative comparison of different repeats.

### Base-level understanding of crossover and non-crossover recombination events in *msh1*

We investigated the internal single-nucleotide variants (SNVs) and indels for the 27 groups of high- and intermediate-activity repeats that were not 100% identical in sequence (i.e., after excluding the perfect repeats Large1, Large2, C, F, K, L, Q, and Z). Conducting such a comprehensive analysis at a single-molecule level is not possible with short-read sequencing. We first examined the distribution of crossover sites in one-rearrangement chimeric reads based on the SNVs/indels that distinguish repeat homologs. We found that recombination breakpoints were most likely to occur in the longest identical fragment (LIF) within two-copy repeats and that the proportion of supporting reads was positively correlated with LIF length as a fraction of total repeat length (Figure 4a; Figure S8; Table S17). This indicates that recombination breakpoints are distributed relatively evenly across the length of repeats rather than being localized to a single hotspot.

**Figure 4.**
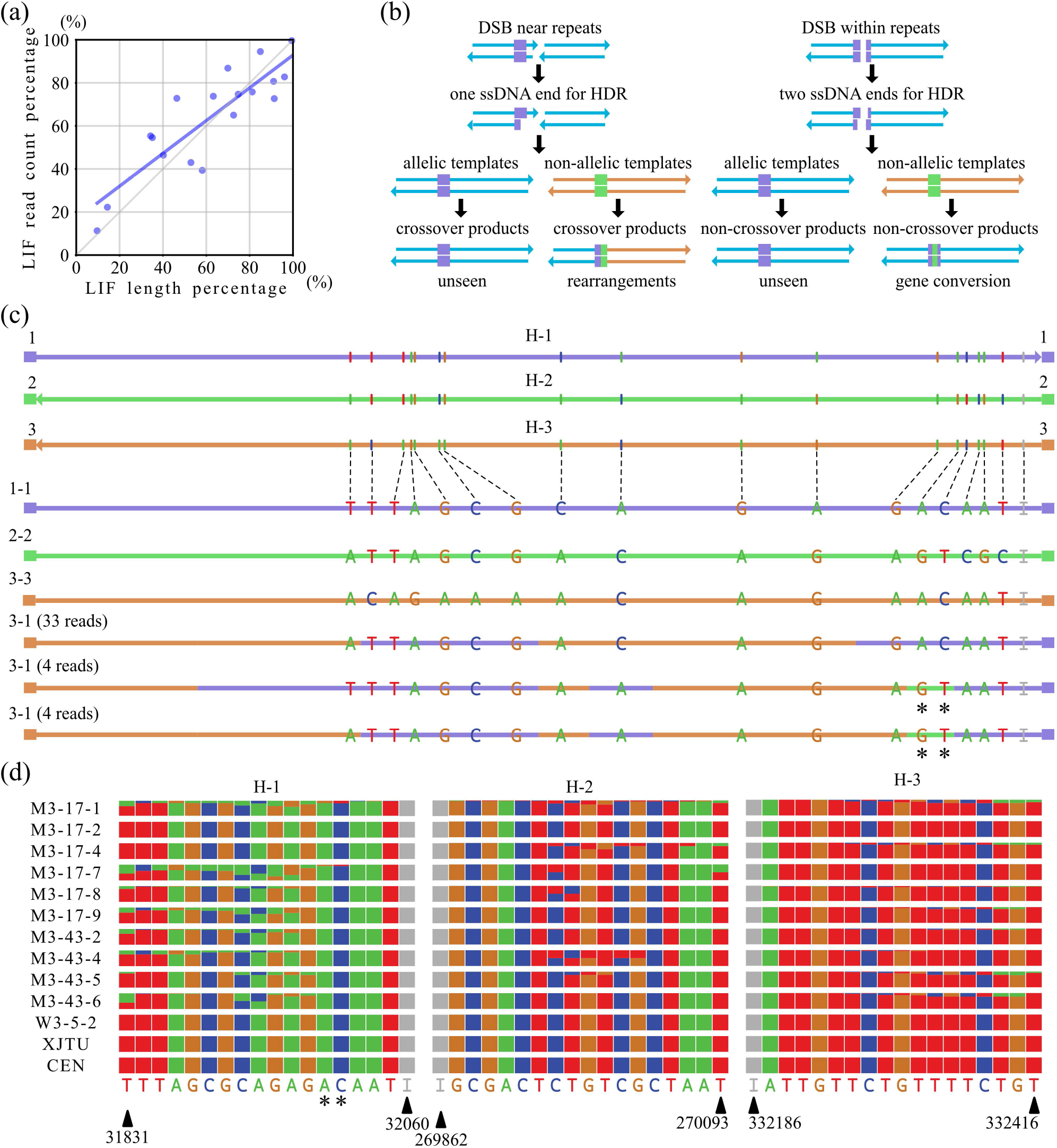
Exchange of SNVs/indels between imperfect repeats is suppressed by *MSH1* (a) Blue circles denote data points from each group of two-copy imperfect repeats. The x coordinate represents the length of the longest identical fragment (LIF) as a percentage of the total length of the repeat. The y coordinate represents the corresponding percentage of recombinant reads for that repeat pair. Blue line denotes the linear regression line. The diagonal 1:1 line denotes equal ratios. Double-strand breaks (DSBs) in DNA can be repaired by allelic or non-allelic dsDNA templates. The purple and green boxes are a pair of imperfect repeats that mediate ectopic homology-directed repair (HDR). The events are not detected (unseen) if allelic templates are involved. Rearrangements or gene conversion events involving non-allelic templates can be detected by sequencing. (c) Discontinuous patterns of SNVs/indels in a proportion of one-rearrangement chimeric reads of repeat H-1 and H-3 may indicate complex patterns of exchange in *msh1* mutants (data from more reads found according to Table S9). The colors of horizontal bars indicate the most parsimonious explanations for the SNVs/indels according to repeat flanking regions. Asterisks denote SNVs potentially introduced by gene conversion from repeat homolog H-2 in these H-1/H-3 mediated one-rearrangement reads. Numbers in parentheses denote the sum of read counts in all samples without normalization. The top three lines in panel c are to scale with vertical bars indicating bases of the same colors as the letters. The lines below are not drawn to scale to avoid crowding. Vertical bars in panels a and c denote A (green), T (red), G (brown), C (blue) for SNVs and I (gray) for indel sites. (d) Square-shaped stacked bar plots are proportions of non-allelic SNVs/indels at predicted sites in repeat group H, which illustrate unsuppressed accumulation in *msh1* mutant samples compared to wild-type (W3-5-2, this study; XJTU, Wang et al., 2021; CEN, Naish et al., 2021). The relative heights in each square-shaped box indicate the ratios of given bases in zero-rearrangement reads for A (green), T (red), G (brown), C (blue), and I for reference (gray) or non-allelic (purple) sequences at indel sites. Reference bases are indicated below. SNVs and indels are arranged from small to large coordinates in the reference. Coordinates of the first and last SNVs/indels (black triangles) are shown. H-2 and H-3 are in the reverse orientation compared to H-1. Asterisks denote the same labeled positions as in panel c.

An earlier study also observed the continuous distribution of SNVs/indels in imperfect repeats I, R and X (Miller-Messmer et al 2012). In our analysis, we also identified discontinuous patterns of SNVs/indels across the length of imperfect repeats in some reads (Table S17). For example, a proportion of the one-rearrangement reads mediated by repeats H-1 and H-3 also contained variants that derived from repeat H-2, suggesting that discontinuous SNV patterns could be related to other repeat homologs not directly involved in a crossover recombination event (Figure 4c). A previous study used the term “partial gene conversion” and suggested mismatch repair mechanism independent of *MSH1* as a possible explanation for discontinuous SNV/indel patterns (Davila et al., 2011).

The patterns of mt genome recombination indicate that dsDNA templates for DSB repair can be allelic or non-allelic, and recombination products can be both crossover and non-crossover (Figure 4b). HiFi technology enabled unambiguous mapping of reads that did not contain any rearrangements but still exhibited variants within repeats (Figure S9). We investigated the exchange of non-allelic SNVs/indels in zero-rearrangement reads spanning the entire length of all 27 high- and intermediate-activity groups of imperfect repeats. We observed numerous mosaic repeat sequences apparently generated through exchange of non-allelic SNVs/indels in *msh1* mutants, which seldomly occurred in wild-type (Table S18-S20). This suggested that without MSH1, non-allelic dsDNA templates were used in non-crossover recombination. In *msh1* mutants, repeat groups A, D, G, H, I, M, MMJS, N, R, T, and W showed a high frequency of non-allelic SNVs/indels (Figure 4d and S10) and had higher frequencies of crossover recombination (Table S16), while the remaining 15 analyzed repeat groups showed low frequencies of non-allelic SNVs/indels (Table S18; note that E-1 and E-2 are not included in this analysis because the same SNV/indel positions are included within the H repeat group). It appears that exchanges can involve very short stretches because the ratios of variants at adjacent positions often differ.

The discontinuous patterns of SNVs/indels in one-rearrangement reads such as the examples described for H-1 and H-3 above could also result from the recombination products of zero-rearrangement templates that have previously undergone these internal exchanges. To investigate this possibility, we compared samples with high (M3-17-1, 16.2%) and low (M3-43-2, 0%) ratios of exchanged zero-rearrangement reads at two positions (the asterisk-labeled A and C in Figure 4d). As predicted, we found the proportion of crossover products between repeat H-1 and H-3 that contained the H-2-specific SNVs (G and T) was also higher in sample M3-17-1 (20.1%) than M3-43-2 (2.0%). The ratios of the same variants in repeat group H were different among *msh1* individuals, which might reflect the process of heteroplasmic sorting (Figure 4d and S11). But in repeat homolog MMJS-2, the unusually high ratios of non-allelic indels in some individuals (M3-17-4 and M3-43-6) were associated with the near complete loss of parental forms, i.e., very low sequence coverage in this region (Figure S10 and S12). Together, these sequence-level investigations suggest that *MSH1* can suppresses the exchange of non-allelic SNVs/indels between imperfect repeats within the mt genome in addition to suppressing structural variation.

### Mt genome evolution in Arabidopsis involves reproducible sets of repeats

We used metaFlye to reassemble the mt genomes of seven other *Arabidopsis thaliana* accessions using published PacBio continuous long-read (CLR) datasets (Jiao and Schneeberger, 2020). We found that all the metaFlye assembly graphs were (near) completely resolved, which suggests low heteroplasmy in all accessions (Figure S13; Table S21 and S22). In contrast, the graphs were much more complex with fragmented contigs and many unresolved repeats in *msh1* mutants, reflecting highly rearranged and heteroplasmic mt genomes (Figure S14a). The graphs of wild-type Col-0 based on published HiFi datasets (Wang et al., 2021; Naish et al., 2021) were still relatively simple (Figure S14b), which indicated that the complexity of mt genomes of *msh1* mutants was not caused by higher sequence depth.

Even though the in vivo physical structures of mtDNA molecules are mostly branched linear forms and circular molecules much smaller than the expected size of a complete master circle (Bendich et al., 1993; Kozik et al., 2019), the “master circle” models facilitate easier comparison and clearer presentation of structural variants. When comparing the pseudo-master circles of multiple accessions with Col-0 mt genome, we identified structural rearrangements associated with specific non-tandem repeats, MMEJ/NHEJ and tandem repeats, as well as some large indels flanked by repeats or microhomologies (Figure S15; Table S23 and S24). We noticed that the repeats that mediated rearrangements, and the repeats or microhomologies flanking the large insertions were often shared by different accessions (Figure S15; Table S24). For example, the structures of mt genomes were strikingly similar in An-1 and L*er*. The mt genomes of Col-0 and An-1 differ by five rearrangements mediated by repeat group B, I, H-1/2, L and H-1/3, while that of L*er* has an additional rearrangement mediated by repeat group LL (Figure 5a). The small circles generated by closely located direct repeats (H-1/2, H-1/3 and LL) were lost during evolution, but in certain cases they can behave like autonomously replicating episomes (Wallet et al 2015; Chevigny et al, 2022). Despite this apparent reproducibility, there were also examples of accession-specific large indels, including a 1105-bp sequence absent Col-0, a 1790-bp sequence absent in C24, a 981-bp sequence absent in Kyoto, and a 1407-bp sequence uniquely present in Sha. The 1407-bp sequence in Sha is flanked by an 18-bp microhomology and QQ-2 sequence and consists of a 1025-bp large repeat plus a 400-bp sequence of unknown origin, which is thought to be associated with the CMS phenotype of Sha (Gobron et al., 2013). Interestingly, we observed that the 1025-bp large repeat in Sha was flanked by a pair of the same 18-bp microhomologies, which occurs 13 times in Col-0 mt genome. Thus, large repeats in the mt genome may simply originate by a two-step “excision-reinsertion” mechanism, in which the excision step is an MMEJ event mediated by a pair of microhomologies and the reinsertion step involves a third microhomology in an unexcised copy of the genome (Figure 5b).

**Figure 5.**
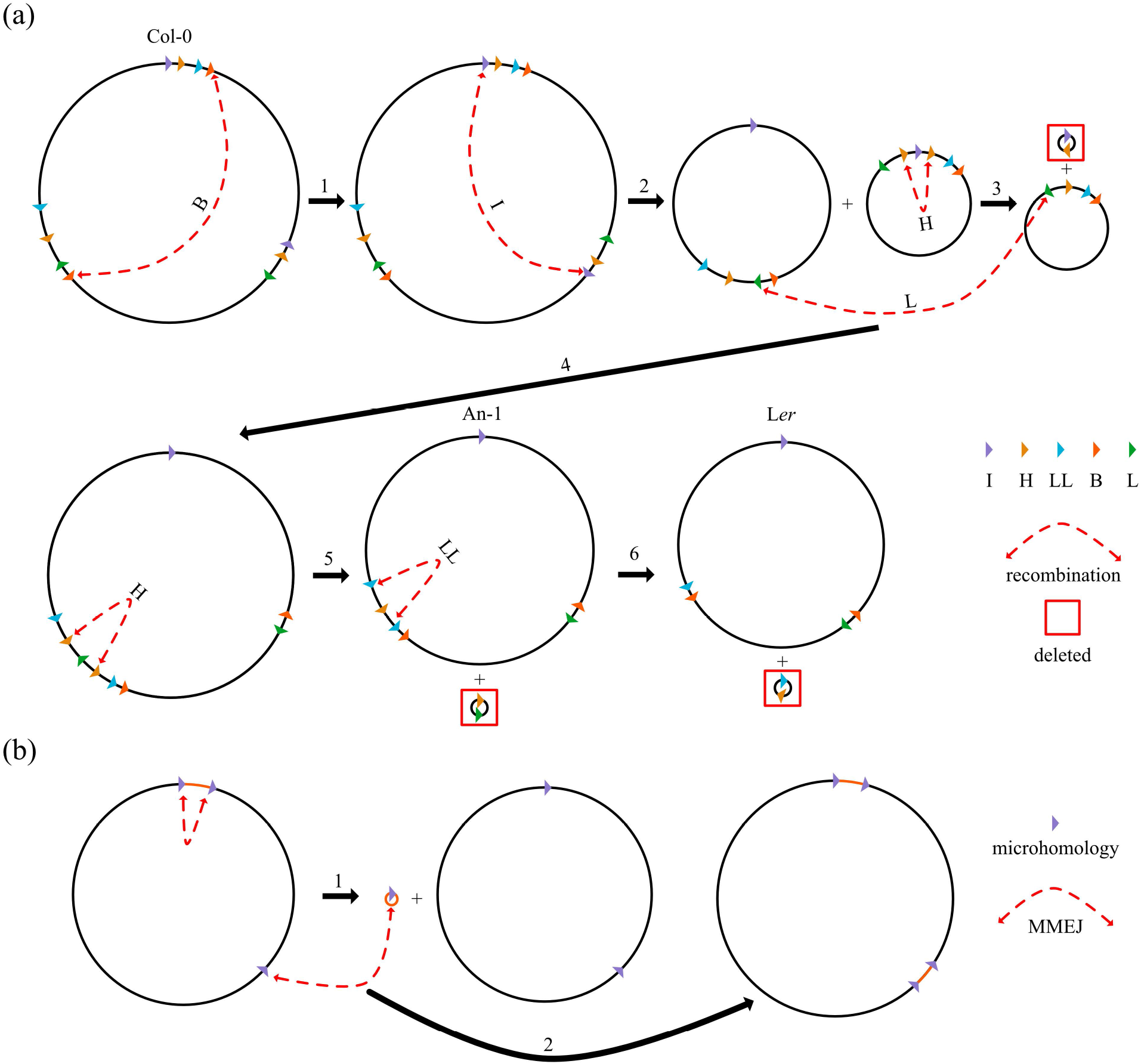
Proposed models for the generation of new mt genome configurations and large repeats (a) Five rearrangements can explain the structural differences and three large deletions (1002-bp, 15652-bp, 55-bp) that distinguish the Col-0 and An-1 mt genomes. An additional rearrangement is needed to explain the large deletions (5920-bp, 15652-bp, 944-bp) in L*er*. Note that all three copies of repeat group H are involved. The 1105-bp sequence absent in Col-0 was not considered in this model. (b) A two-step “excision-reinsertion” mechanism for the generation of large repeats in mt genomes. Note that microhomologies with three copies (or more) are needed.

The overall HiFi read coverage of different *msh1* individuals did not vary greatly compared with wild-type, except for fluctuations at sporadic stretches (Figure S16). The region around repeat Large1-2 showed the lowest coverage in all *msh1* individuals, which would indicate a partial loss of *trnS*, *atp6-2* and *atp9* (Figure S17a-c). By analyzing the depth of CLR reads, we verified the loss of two-copy genes *trnS* and *atp6-2*, but not the single-copy gene *atp9* (Figure S17d-f). Together, our analyses suggest that both repeat-mediated rearrangements and the relatively random MMEJ/NHEJ observed in *msh1* mutants are involved in mt genome evolution, leading to rearrangements, large deletions, and generation of new repeats with selection preventing the loss of essential functional genes.

### Different accumulation patterns of pt genome variants in *msh1* mutants

The frequencies of structural variants were very low in the pt genome for both wild-type and *msh1* mutants (Figure S18; Table S25-S27). In contrast to the mt genome, the Arabidopsis pt genome contains a pair of very large repeats (IRa/b) but very few intermediate repeats (Table S28). We detected only one read in a single sample that supported the previously identified rearrangement mediated by a pair of 123-bp imperfect repeats (Xu et al., 2011). Therefore, this variant appears to make a very small contribution towards pt genome instability. In *msh1* mutants, the proportion of pt MMEJ events was increased as also observed in mt genomes, but the associated microhomologies included an overrepresentation of short, partial palindromic sequences (Figure S18; Table S29). A partial palindromic sequence can be viewed as a pair of repeats or microhomologies with both copies at the same location but in inverted orientation. For example, in MMEJ events mediated by the most active partial palindromic sequence (a 22-bp stem and a 9-nt loop) in *msh1* mutants, we observed recombination breakpoints in both the stem and the loop region, resembling observations in mt genomic imperfect repeats (Figure 6a; Table S30).

**Figure 6.**
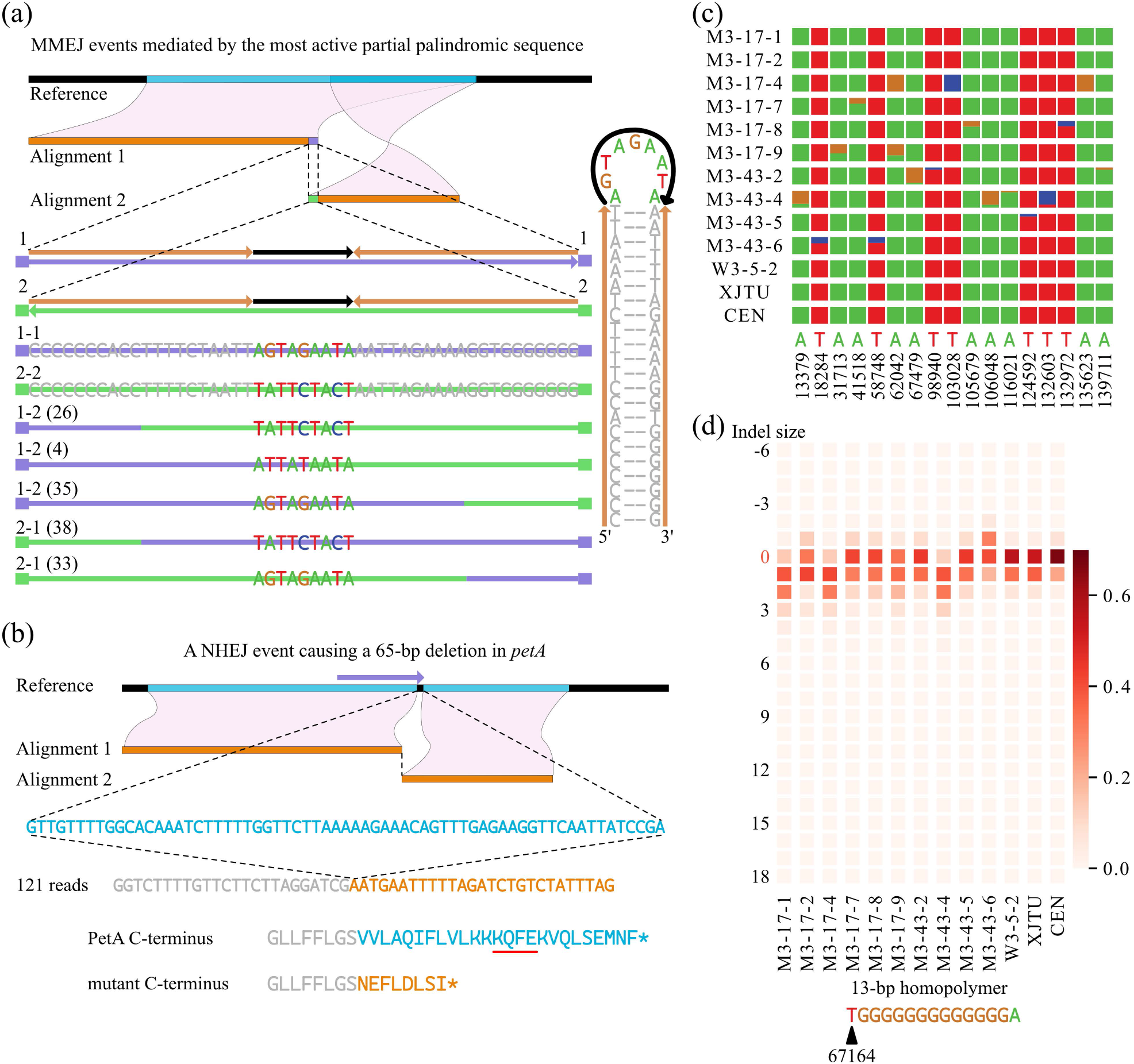
Mutations and structural variants in the pt genome detected by HiFi reads (a) The partial palindromic sequence contains a stem (orange arrow) and a loop (black arrow) and is also a pair of repeats or microhomologies with both copies at the same location but in inverted orientation. The original sequence can be viewed as repeat homolog 1 (1-1), and at the same time as repeat homolog 2 (2-2) in the opposite direction. MMEJ events mediated by the most active partial palindromic sequence in *msh1* mutants were shown. Numbers in brackets denote the sum of read counts showing crossover site distribution in chimeric reads in all samples without normalization. Note that there are four reads showing the breakpoint in the middle of the loop. The ratio of one-rearrangement reads versus zero-rearrangement reads across this partial palindromic sequence is 160/14786 (1.1% of total reads) in *msh1* and 0/1966 in wild-type. (b) A NHEJ event occurring only in the M3-43-5 individual was predicted to disrupt the *petA* gene. The ratio of one-rearrangement reads versus zero-rearrangement reads across this deletion is 121/1385 (8.0% of total reads). (c) Square-shaped stacked bar plots are proportions of AT-to-GC transitions in 17 GATK-called SNVs in wild-type and *msh1* mutant individuals, which were enriched in one or a few samples at high ratios. The relative heights in each square-shaped box indicate the ratio of a given base for A (green), T (red), G (brown), and C (blue). Reference bases are indicated below. (d) Heatmap shows the proportions of length variants at a 13-bp homopolymer started near position 67164.

Another important feature was the sample-specific accumulation of certain variants in pt genomes of *msh1* mutants. Although most NHEJ products were at low frequencies with a seemingly random distribution, we observed one pt NHEJ product that occurred only in one sample (M3-43-5) but at relatively high frequency (121 one-rearrangement reads versus 1385 zero-rearrangement reads), resulting in a 65-bp deletion in the *petA* gene (Figure 6b; Table S31). This mutation was predicted to delete a highly conserved KQFE motif in the C-terminal stromal extension of cytochrome f, which would compromise photosynthesis (Choquet et al., 2003). This sample-specific accumulation pattern was also observed for SNVs in the pt genome. In total, we identified 17 pt SNVs that were enriched to high frequencies in one or a few *msh1* mutant samples (Figure 6c; Table S32). They were predominantly AT-to-GC transitions, consistent with the mutation spectrum in *msh1* mutants that was reported previously and suggested to be caused by polymerase errors (Wu et al., 2020). GC variants were also recently shown to be favored in the process of heteroplasmic sorting in *Arabidopsis thaliana* (Broz et al., 2022). Moreover, we found that simple sequence repeats (predominantly in homopolymers) exhibited larger proportions of length variants in the pt genomes of *msh1* mutants compared with wild-type (Figure S19; Tables S33). For example, for the 13-bp homopolymer at position 67164 (which had reads supporting length variants ranging from - 6 bp to +18 bp), the proportion of reference length (13 bp) was less in all *msh1* individuals than wild-type (Figure 6d). Together, our sequence data suggest that MSH1 participates in removal of pt genome variants. The specific pt variants in *msh1* mutants are less reproducible among samples than mt variants, and heteroplasmic sorting may contribute to this variation among individuals.

## Discussion

In this work, we used HiFi reads (~15 kb long, > 99% accuracy) to study the patterns of mt and pt genome variants using samples from Arabidopsis *msh1* mutant individuals. HiFi reads were long enough to span the entire lengths of repeats, including the two largest repeats in the Arabidopsis mt genome (6.5 kb and 4.2 kb), which enabled analyses of recombination breakpoints and higher-order structural rearrangements that were not previously possible with short reads. We developed a mapping-based pipeline, which showed that HiFi reads performed well in resolving one to multiple repeats-mediated rearrangements in a single read and studying polymorphisms between long non-identical repeats. Due to the ancient conservation of the *MSH1* gene across angiosperms and green plants in general (Wu et al., 2020), our analysis in Arabidopsis likely extends to mt genomes in other plants. Compared to previous studies, we also identified additional repeats active in low-frequency recombination. Similarly, in the bacterial *Borrelia burgdorferi*, long-read sequencing successfully identified a 97% reduction in the activity of recombinational switching at the *vlsE* locus in MutL-deficient line, which was much higher than the ratio (36%) estimated by short-read sequencing (Castellanos et al., 2022). However, due to the low read counts, HiFi reads were less cost-effective for studying low-frequency MMEJ and NHEJ variants. This cost could be reduced by isolating organelles from the cells before DNA extraction for sequencing.

### MSH1 suppresses exchange of SNVs and indels between non-identical repeats

There might be a common need to suppress recombination mediated by non-allelic genomic repeats across systems. During meiosis in the nucleus, MutS homologs detect mismatches and recruit the DSB-generating SPO11 complex to suppress crossover repair between non-identical centromeric repeats (Blackwell et al., 2020; Naish et al., 2021). In the mitochondria, MSH1 is similarly required to suppress crossover repair between intermediate-sized repeats (Arrieta-Montiel et al., 2009; Davila et al., 2011). Gene conversion was previously proposed to explain the discontinuous pattern of SNVs/indels derived from non-identical repeats in chimeric reads (Davila et al., 2011). However, the short-read sequencing performed in that study makes it difficult to assess whether these discontinuous patterns were present even in molecules that did not exhibit structural rearrangements from recombination. In our analysis, unambiguous mapping of HiFi reads across the full length of mt repeats identified the exchange of SNVs and indels between pairs of imperfect repeats in zero-rearrangement reads in *msh1* mutants. MSH1 activity appears to suppress these exchanges based on their relative absence in wild-type.

The observed variants could be introduced by non-crossover gene conversion events, but they could also result from multiple rounds of crossover recombination involving the same repeats but different breakpoints. It is important to note that any analysis of a standing population of genome copies in a tissue sample will not only reflect the recombination events that originally generated structural rearrangements but also the subsequent history replication, degradation, and heteroplasmic sorting. Furthermore, such processes may be sensitive to rearing conditions (e.g., day length) and the stage in development when tissue was harvested, which should be considered in any comparison between our results and previous studies.

There could very well be a connection between MSH1’s functions in suppressing ectopic recombination and mismatch repair mechanisms. Specifically, MSH1 may provide a surveillance mechanism to check for mismatches in double-stranded DNA regardless of whether they are introduced by polymerase errors or by strand invasion between short or non-identical repeats, followed by introduction of DSBs to (re)start repair by homologous recombination. To suppress recombination at short and intermediate repeats that are 100% identical in sequence, MSH1 may need to detect differences in boundary sequence. During homologous recombination, the bacterial RecA-ssDNA presynaptic filament checks homology in ~9 bp steps before strand exchange with the dsDNA template (Huang et al., 2020). In such screening mechanisms, the first mismatches may be detected at the repeat boundaries when the repeat sequences themselves are identical. Thus, MSH1 may remove both crossover and non-crossover forms of ectopic recombination to ensure accurate DSB repair of the genome by detecting mismatches within repeats or at repeat boundaries (Figure 7a).

**Figure 7.**
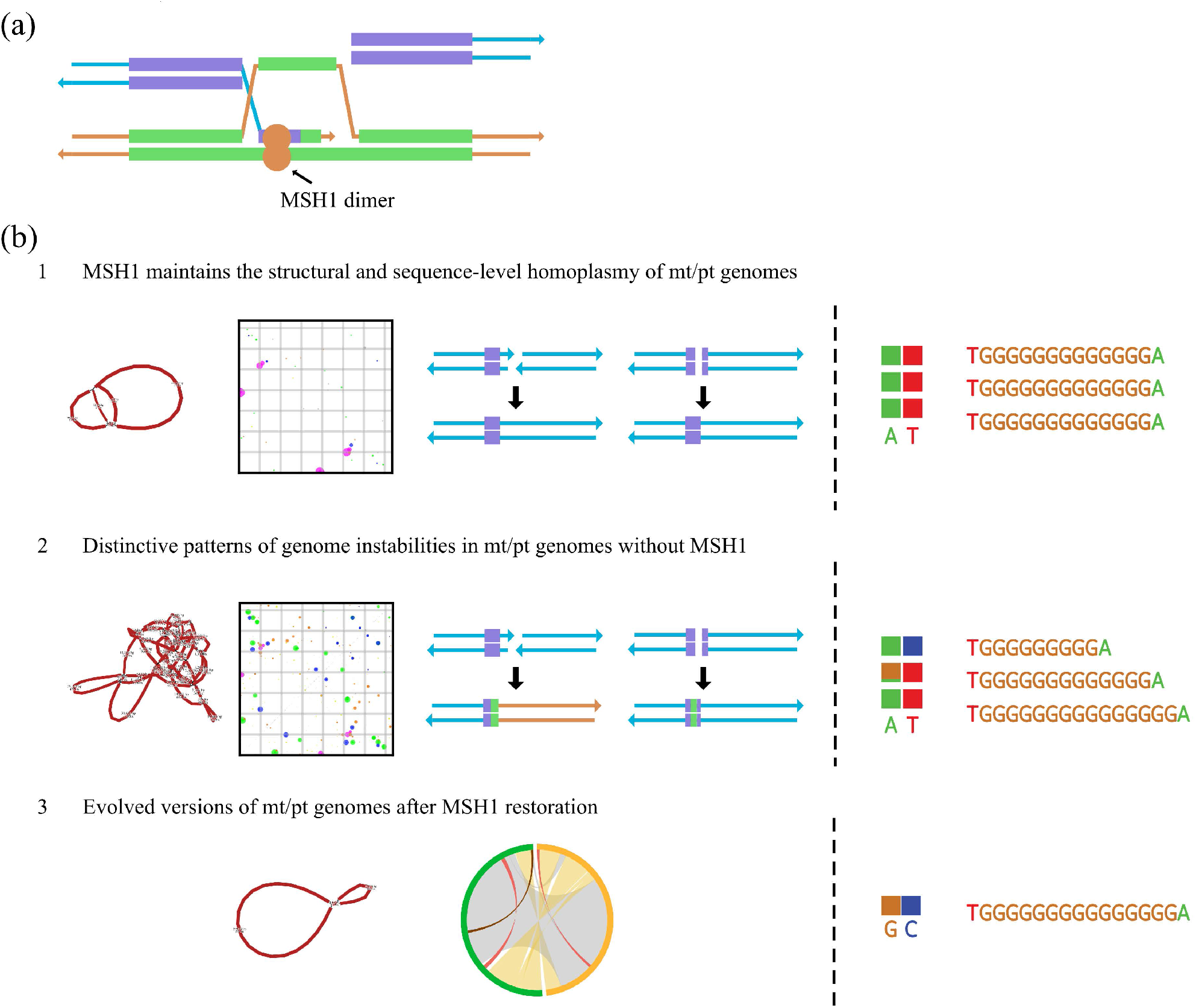
Proposed models of mt/pt genome evolution controlled by fluctuating *MSH1* levels (a) MSH1 appears to suppress both crossover and non-crossover forms of ectopic recombination by detecting mismatches within repeats or at repeat boundaries if the two copies of repeats are identical. (b) In the wild-type (Step 1), MSH1 maintains the structural and sequence-level homoplasmy by repressing ectopic recombination (and by removing other types of variants resulting from polymerase misincorporations and/or DNA damage). It has been previously hypothesized that MSH1 performs these functions by recognizing mismatches in DNA and introducing DSBs to (re)initiate recombinational repair. When *MSH1* is disrupted either genetically or environmentally (Step 2), organellar genomes become highly heteroplasmic. When MSH1 activity is restored (Step 3), both genomes return to a low-homoplasmy state. The new mt genome undergoes rearrangements and large deletions involving reproducible sets of repeats. Given the low abundance of intermediate-sized repeats in Arabidopsis pt genome, altered pt genome often differ at random sites relating to MMEJ/NHEJ, AT-to-GC mutation, or length variants at simple sequence repeats.

### Conventional recombination activity only accounts for a fraction of DSB repair events

Most previous mt repeat annotations depended on the penalty parameters of mismatches and gaps used in sequence alignment. We generated a global and quantitative map of rearrangements mediated by repeated sequence elements, including the identification of additional low-frequency repeats, which raises new questions about what contributes to the observed differences in recombination activity among repeat pairs. Increased crossover recombination events involving intermediate-sized repeats have been observed in wild-type plants when DSB frequencies were increased by DNA damaging chemicals (Wallet et al., 2015). Therefore, repeats near DSB hotspots may show higher ectopic recombination activity compared to other repeats with similar length and sequence identity, which might explain cases that fall far from the regression line that defines the relationship between recombination activity and repeat length (Figure S5b). It may also help to study repair events systematically by inducing DSBs through mt-targeting CRISPR-Cas9 or mitoTALENs (Arimura et al., 2020) and in more RRR gene mutants, such as *why2*, *reca3*, *osb1* (Gualberto and Newton 2017).

DSBs within a repeat or in its flanking sequences would be repaired using homologous DNA stretches of another intra- or intermolecular copy. After resection of DSB ends, RecA recombinases bind 3’ ssDNA and rapidly search for homology (Wiktor et al., 2021). It was reported that at least 8-nt microhomology was needed for RecA-ssDNA filaments to bind dsDNA templates stably, and 15-nt was sufficient to be resistant to facilitated exchange during homology search (Qi et al., 2015). These studies suggest that only a small stretch of RecA-ssDNA is needed during each HDR repair event. Typically, only crossover products have been used to estimate frequencies of recombinant conformations (Wallet et al., 2015; Odahara et al., 2017). Such calculations do not account for the fact that non-crossover recombination or gene conversion events between repeats can also occur at substantial frequencies. Because *msh1* mutants exhibited exchange of sequence variants between non-identical repeats even in reads with no structural rearrangements, the increases in ectopic recombination activity are likely underestimated when only crossover products are quantified.

When calculating recombination for some three-copy or four-copy repeats locating within other longer two-copy repeats, one-rearrangement reads shared by both shorter and longer annotations were assigned only to the larger two-copy repeats by blastn (Figure 3). This illustrates that it may be more natural to view recombination events as associated with specific genomic positions, but not with repeat annotations.

### Different variant accumulation patterns in mt and pt genomes

In plant organellar genomes, large repeats recombine constantly. The numerous intermediate-sized repeats are the major source of altered recombination activities in *msh1* mutants. The global patterns of mt recombination indicated by one-rearrangement reads (Figure 2) and fluctuation in sequence depth (Figure S16) were similar among individuals, which could be a combined effect of the locations of repeat pairs and the selection to maintain regions containing functional genes. Despite the common loss of the region around repeat Large1-2 (containing *atp6-2*, *atp9* and *trnS*), the relative dosage of parental and crossover recombination forms of each repeat pair may differ among *msh1* individuals and generations, which was proposed to be caused by heteroplasmic sorting in previous studies (Shedge et al., 2007; Arrieta-Montiel et al., 2009). In other Arabidopsis accessions, we observed large indels flanked by repeats or microhomologies. These indels sometimes coincided with the low-coverage regions observed in *msh1* mutants, including the previously investigated region around Large1-2. Some of these structural rearrangements may cause a CMS phenotype, such as the one reported in Sha (Figure S15; Gobron et al., 2013). In natural populations, the mt genome may be transiently (within a single generation) found in a highly rearranged and heteroplasmic states similar to *msh1* mutants, providing source materials for functional innovation (Woloszynska, 2010; Tang et al., 2017; Johnston, 2019). However, the metaFlye assembly graphs support previous observations that the mt genomes of wild-type Arabidopsis accessions typically exist in low-heteroplasmy states with structural variants predominantly associated with repeats or microhomologies mediated DSB repair activities (Figure S13; Table S24).

The contrasting patterns of variation between mt and pt genomes may result from differences in repeat spectrum, genome copy number per organelle, and DNA repair mechanisms. Structural rearrangements in pt genome of Arabidopsis *msh1* mutants were hardly detected in earlier studies, possibly due to the lack of intermediate-sized repeats (Xu et al., 2011). HiFi-identified rearrangements in the pt genome were mostly MMEJ and NHEJ events. The MMEJ events mediated by partial palindromic sequences were found at a higher frequency than others and showed repeatability among samples. Moreover, other examples of pt genome variants occurred at high frequency in individual samples regardless of whether they were MMEJ, NHEJ (Figure 6b) or SNVs (Figure 6c). Such sample-specific variants were most likely due to a combination of differential occurrence of the original mutations and subsequent variation introduced by heteroplasmic sorting. Simple sequence repeats including homopolymers are intrinsically unstable regions in pt genomes, which is also described in an earlier characterization of the mutation spectrum of 51 *Oenothera* mutants (Massouh et al., 2016). Although length variants at these homopolymers are found even in wild-type plants, our data suggest that the frequencies of these variants are reduced by *MSH1* activity (Figure 6d and S19).

In conclusion, *MSH1* acts to maintain both structural and non-structural homoplasmy in plant organellar genomes possibly by promoting gene conversion (with allelic DNA templates). These genomes can be transiently disrupted when *MSH1* expression is downregulated, providing more raw materials for further functional selection (Yang and Mackenzie, 2019). When *MSH1* expression is restored, evolved versions of the genomes can rapidly rise in frequency (Shedge et al., 2007). However, the two genomes can respond quite differently to the loss of the same gene (Figure 7b). Organellar genome editing technologies are becoming more and more accessible (Kazama et al., 2019; Arimura et al., 2020; Forner et al., 2022). However, most protocols in non-model species and the recent TALEN-GDM in Arabidopsis require a callus-induction step (Forner et al., 2022), during which the expression of MSH1 may be downregulated (Sun et al., 2012). Thus, we need to pay attention to the rearrangements and gene conversion events in mt genomes, which are not easily detected by short-read sequencing, as well as the unwanted random accumulation of AT-to-GC mutations and length variants of simple sequence repeats in pt genomes. In the future, we may better understand the role of cytoplasmic genetic material in crop improvement through studying RRR genes including *MSH1*.

## Methods

### DNA extraction and PacBio HiFi sequencing

Generation of *msh1* mutant and wild-type F_3_ families of *Arabidopsis thaliana* was described previously (Wu et al., 2020). In brief, an *msh1* mutant (*chm1-1*, CS3372) was used as the pollen donor in a backcross to *A. thaliana* Col-0 so that the F1 plant would have wild-type organellar genomes. Seeds of each homozygous mutant and wild-type F2 plant was collected separately and propagated as F_3_ families. To extend rosette stage growth before bolting, plants were grown in a short-day growth chamber (8 h day and 16 h night) for six weeks. Rosettes of 11 individuals were collected to extract DNA with the CTAB method. Total DNA was purified with a QIAGEN kit (Q13343) and quantified using NanoDrop and Qubit. Purified DNA (3.3~18.6 μg for each sample) was used for the construction of SMRTbell libraries according to PacBio’s standard protocol. Sequencing was performed using the PacBio Sequel II platform in CCS mode, which generated a total of 34.36 Gb HiFi reads (12.7~18.1 kb mean read length). Both the mutant and wild-type individuals used in our studied may have segments originated from the L*er* accession in the nuclear genome as indicated by an earlier study (Virdi et al., 2015).

### Identification of rearrangements in mt and pt genomes

Raw HiFi reads from each sample were mapped to the Arabidopsis Col-0 reference genome (TAIR10 for nucleus; NC_037304.1 for mitochondria; NC_000932.1 plus a 1-bp expansion at position 28673 in the pt genome as described in Wu et al., 2020) with minimap2 (version 2.20, r1061; parameters: -ax map-hifi; Li, 2018). Reads mapped to mt and pt genomes were separated using bamtools (version 2.5.1; Barnett et al., 2011) and realigned against each genome with blastn (version 2.12.0; Camacho et al., 2009). Raw blastn results were further processed by a custom pipeline (available at https://github.com/zouyinstein/hifisr) to group the reads by discontinuity, alignment overlap (AO) size (i.e., repeat size) and mapping coordinates, and calculate read counts for input samples. Different types of genome rearrangements were manually checked, including zero-, one-, two-, three-, and four-rearrangement reads, MMEJ (AO length of 2-49 bp), NHEJ (AO length of 0-1 bp), and replication slippage mediated by tandem repeats. We also identified large insertions, which were poly-G/C tracts in most cases. However, it was not clear whether these inserted homopolymers were biological or a sequencing artefact, and they were not analyzed further.

### Mitochondrial repeat annotations and recombination activity

Candidate coordinates of repeats were generated by blastn (version 2.12.0+; parameters: - word_size 4) using the mt reference sequence as both query and subject (Table S10). The resulting coordinates were manually checked against the junction coordinates of one-rearrangement HiFi reads. We identified 71 groups of two-copy repeats, 23 groups of three-copy repeats, 5 groups of four-copy repeats, which accounted for most junctions of read alignments in *msh1* mutant mt genomes (Table S11-S13). These 99 repeat groups contained all previously reported active repeats except for repeat TT and 53 additional groups of recombinationally active repeats ≥ 50 bp, which we named HNR1 to HNR53 (repeats FF, WW, XX, YY, and ZZ were not considered because they are absent in the NC_037304.1 annotation). For consistency with blastn alignments, the boundaries of some low-identity repeats in our annotation differed from previous annotations. For example, the direct repeats MMJS-1 (767 bp) and MMJS-2 (692 bp) were previously annotated as separate smaller repeats MM, J, and S. We treated short repeats as part of the microhomology category (2 to 49 bp) because blastn can often generate overlapping, inaccurate, or incomplete predictions of homology below 50 bp in size. Per base ratios of rearranged reads versus total reads and read counts for parental and recombinant forms were calculated with custom Python scripts based on blastn outputs of the HiFi-SR pipeline and used to identify dynamic regions of the mt genome.

### Identification of non-allelic SNVs/indels between imperfect repeats

To analyze patterns of SNVs/indels, zero-rearrangement reads of each sample were re-aligned to the mt reference genome by PacBio pbmm2 (version 1.3.0; parameters: --sort --preset CCS). The resulting BAM files were used for variant calling by GATK HaplotypeCaller (version 4.2.3.0). Variant frequencies and sequencing depths for each GATK-called SNV/indel were calculated from raw BAM files with the Python module pysam (version 0.17.0). Non-allelic SNVs/indels inferred from putative gene conversion events between some high-activity mt imperfect repeats were not always detected by GATK but were also used for calculations.

### Calculation of absolute and normalized per base coverage

Total mt genome-mapped reads were re-aligned to mt reference genomes by PacBio pbmm2 (version 1.3.0; parameters: --sort --preset CCS). Absolute per base coverage across the entire mt genome of each individual was calculated with the samtools depth subcommand (version 1.7) and normalized to total bases of each sample. Annotations are based on the GenBank file NC_037304.1 (Sloan et al., 2018).

### metaFlye assembly and synteny analysis of mt genomes

Raw PacBio CLR reads from each accession were mapped to the reference genomes (the respective nuclear assembly in Jiao and Schneeberger, 2020; Col-0 NC_037304.1 for mitochondria and NC_000932.1 plus a 1-bp expansion at position 28673 for plastid) with minimap2 (version 2.20, r1061; parameters: -ax map-pb; Li, 2018). Reads mapped to mt genomes were separated using bamtools (version 2.5.1; Barnett et al., 2011) and were re-assembled with metaFlye (Flye version 2.9-b1774; parameters: --meta --pacbio-hifi/--pacbio-corr --extra-params output_gfa_before_rr=1 --genome-size 370K/2M for mitochondria; Kolmogorov et al., 2020). Before assembling, PacBio CLR reads longer than 1000 bp were extracted by Filtlong (version 0.2.1; parameters: --min_length 1000) and were corrected using CONSENT (version 2.2.2; parameters: --type PB; Morisse et al., 2021). Assembly graphs before and after repeat resolution were visualized by Bandage (version 0.8.1; Wick et al., 2015). Reads from different *msh1* families were merged. To get linear representations of pseudo-master circles for mapping-based analysis, we started from the assemblies (--genome-size 2M) after repeat resolution, discarded low-coverage contigs, manually resolved the large repeats for accessions Eri-1, C24, Cvi-0, An-1, L*er*, Sha, and re-resolved Large2 repeats for Kyoto with the Bandage software, followed by rotation or reverse complementation. Corrected CLR reads were realigned against each pseudo-master circle using the HiFi-SR pipeline. To analyze the shared synteny between mt genomes, we performed blastn alignments using the Col-0 mt reference sequence as the query and pseudo-master circle of other accessions as the subject. Raw blastn results were processed by a custom Python script available in HiFi-SR. Coordinates of large repeats were determined based on blastn alignments using the pseudo-master circle as both query and subject. Repeat-mediated rearrangements, MMEJ/NHEJ events, and large insertions composed of short fragments of multiple origins were manually checked. Plots were generated by the Python package pycircos (version 1.0.2).

### Identification of SNVs/indels in the pt genome

To analyze patterns of SNVs/indels in the pt genome, total reads of each sample were re-aligned to the pt reference genome by PacBio pbmm2 (version 1.3.0; parameters: --sort --preset CCS). The resulting BAM files were used for variant calling by GATK HaplotypeCaller (version 4.2.3.0). Variant frequencies and sequencing depths for each GATK-called SNV/indel were calculated from raw BAM files with the Python module pysam (version 0.17.0).

## Acknowledgments

We thank Xiaojing Zhang from Prof. Li-Zhen Tao’s laboratory at South China Agricultural University for assistance with plant growth. This work was supported by grants from National Natural Science Foundation of China (32170238), the Science, Technology and Innovation Commission of Shenzhen Municipality (RCYX20200714114538196), the U.S. National Institutes of Health (R01 GM118046), and the Chinese Postdoctoral Science Foundation (2021M693451).

## Author contributions

ZW designed the research; YZ performed the experiments; YZ wrote the computer scripts and performed majority of the bioinformatic analysis; WZ analyzed the boundaries of structure rearrangements in Arabidopsis accessions; YZ, WZ, DBS, and ZW discussed the results; YZ, DBS and ZW wrote the manuscript; DBS and ZW revised the manuscript.

## Data Availability

The raw sequence data reported in this paper have been deposited in the Genome Sequence Archive (Chen et al, 2021) in the National Genomics Data Center (C.-N. Members and Partners, 2022), China National Center for Bioinformation / Beijing Institute of Genomics, Chinese Academy of Sciences (GSA: CRA006060), which is publicly accessible at https://ngdc.cncb.ac.cn/gsa. The Python-based pipeline for the detection of plant mt structural rearrangements used in this study is available at https://github.com/zouyinstein/hifisr.

## Supporting Information

**Figure S1** Seedlings of *msh1* mutants and wild-type *Arabidopsis thaliana*

**Figure S2** Mapping ratio of HiFi reads in different samples

**Figure S3** Patterns of one-rearrangement events detected in mt genome of wild-type

**Figure S4** Variability of structural rearrangements across the mt genome

**Figure S5** Comparison of the recombination activities of different repeat groups

**Figure S6** Parental forms and recombination products can be detected according to paths

**Figure S7** Two-rearrangement reads involving large repeats are increased in *msh1* mutants

**Figure S8** Continuous distribution of SNVs in non-identical repeats CC-1/2

**Figure S9** Mapping of zero-rearrangement long reads identifies variants within repeats

**Figure S10** Imperfect repeats with relatively higher ratios of exchange of SNVs and indels in *msh1* mutants

**Figure S11** Coverage of regions containing repeat group H in wild-type and *msh1* individuals

**Figure S12** Coverage of regions containing repeat group MMJS in wild-type and *msh1* individuals

**Figure S13** metaFlye assemblies of mt genomes in other accessions

**Figure S14** The metaFlye assemblies of mt genomes are more complex in *msh1* mutants than wild-type

**Figure S15** Synteny analysis of mt genomes of other *Arabidopsis thaliana* accessions

**Figure S16** Normalized per base read coverage across mt genome

**Figure S17** Coverages around *atp6-1*, *atp6-2*, and *atp9* in *msh1* mutants and Arabidopsis accessions

**Figure S18** The proportions of structural variants in Arabidopsis pt genomes

**Figure S19** Length variants at simple sequence repeats in the pt genomes

**Table S1** DNA content and fresh weight of individual plants used for HiFi sequencing

**Table S2** Mapping statistics of HiFi reads of individual plants

**Table S3** Statistics of mt genome-mapped reads of individual plants

**Table S4** Statistics of pt genome-mapped reads of individual plants

**Table S5** Read counts for different types of mt structural rearrangements in individual plants

**Table S6** Read counts for different categories of mt structural rearrangements in wild-type and *msh1* plants

**Table S7** Alignment overlap length in mt one-rearrangement reads.

**Table S8** HiFi-SR outputs of two-rearrangement mt genome-mapped reads of individual plants

**Table S9** HiFi-SR outputs of one-rearrangement mt genome-mapped reads of individual plants

**Table S10** Raw blastn outputs using the mt reference sequence as both query and subject

**Table S11** Annotations of two-copy repeats based on blastn and one-rearrangement HiFi reads

**Table S12** Annotations of three-copy repeats based on blastn and one-rearrangement HiFi reads

**Table S13** Annotations of four-copy repeats based on blastn and one-rearrangement HiFi reads

**Table S14** Recombination frequencies of two-copy repeats

**Table S15** Recombination frequencies of three-copy repeats

**Table S16** HiFi read counts indicating parental forms and crossover recombination products of representative repeats in wild-type and *msh1* mutants.

**Table S17** Counts of continuous and discontinuous one-rearrangement reads in high- and intermediate-activity two-copy imperfect repeats

**Table S18** Ratios of predicted mt gene conversion sites in individual plants and published HiFi datasets

**Table S19** Frequencies of GATK-identified mt SNVs in individual plants and published HiFi datasets

**Table S20** Frequencies of GATK-identified mt indels in individual plants and published HiFi datasets

**Table S21** Mapping statistics for PacBio CLR reads in individual plants

**Table S22** Statistics of corrected PacBio CLR reads used for metaFlye assembly of mt genomes

**Table S23** Blastn results for mt large repeats

**Table S24** Blastn results for syntenic relationships in mt genomes

**Table S25** Read counts for different types of pt structural rearrangements in individual plants

**Table S26** Read counts for different categories of pt structural rearrangements in wild-type and *msh1* plants

**Table S27** Alignment overlap length in pt one-rearrangement reads

**Table S28** Raw blastn outputs using the pt reference sequence as both query and subject

**Table S29** HiFi-SR outputs of pt one-rearrangement reads mediated by short partial palindromic sequences in individual plants

**Table S30** HiFi-SR outputs of pt one-rearrangement reads mediated by the most active short partial palindromic sequence in individual plants

**Table S31** HiFi-SR outputs of a pt NHEJ event in individual plants

**Table S32** Frequencies of GATK-identified pt SNVs in individual plants and published HiFi datasets

**Table S33** Frequencies of GATK-identified pt SNVs in individual plants and published HiFi datasets

