## Supplemental Figure Legends for "Long-read sequencing characterizes mitochondrial and plastid genome variants in Arabidopsis *msh1* mutants"

**Supplementary Figure Legends**

**Figure S1** Seedlings of *msh1* mutants and wild-type *Arabidopsis thaliana*

(a) Photos show wild-type (W3-5) and *msh1* (M3-17, M3-43) seedlings of different $F_3$ families at 5 days after sowing (DAS). (b) Violin plots show primary root lengths of $F_3$ families at 5 DAS. The shorter root lengths of *msh1* mutant seedlings might indicate delayed seed germination (n = 100; ***, $P < 0.001$; Student's *t* test).

**Figure S2** Mapping ratio of HiFi reads in different samples

Stacked bar plots show ratios of HiFi reads mapped to nuclear (nc), plastid (pt), mitochondrial (mt) genomes by minimap2; un denotes the ratio of unmapped reads.

**Figure S3** Patterns of one-rearrangement events detected in mt genome of wild-type

Dot plot shows global patterns of the junctions and relative read counts of mitochondrial one-rearrangement events in wild-type. Signals from previously named repeats are labeled. Colors represent categories of alignment overlap length (AO length). The x position of each dot is the end coordinate of blastn subject for Alignment 1 (se1). The y position of each dot is the start coordinate of blastn subject for Alignment 2 (ss2). The sizes of dots denote read counts normalized to 10000 total mt genome-mapped reads per sample. The near symmetrical appearance of the plot is a result of HiFi reads aligned in both forward and reverse directions relative to the reference genome. The spots in mirrored positions should not be interpreted as quantifications of the two reciprocal recombination products.

**Figure S4** Variability of structural rearrangements across the mt genome

Ratios of rearranged reads versus total reads are shown across the mt genome in wild-type (W3-5-2) and *msh1* (M3-17 and M3-43). Red horizontal line denotes 50% rearrangements of the mt genome. Grey shaded regions denote highly dynamic regions. Reads from different *msh1* families are merged. Repeat annotations are shown at the bottom (note that annotations of repeats FF, WW, XX, YY, ZZ are not present in GenBank file NC_037304.1).

**Figure S5** Comparison of the recombination activities of different repeat groups

(a) Dot plot shows recombination frequencies of 97 repeat groups (Large1 and Large2 are not included) in *msh1* mutants. Repeat groups that were verified (solid circles) and not verified (hollow circles) to be active in previous studies. Most newly verified active repeat groups belong to the low-activity category with smaller size. (b) Plot shows correlation between homology length and recombination frequency of highly identical two-copy repeats (mismatches + gaps $\leq 2$). Blue line denotes the linear regression line. In (b) and (c), circles above the red dash line belong to the high-activity category (read count $\geq 50$), between the red and purple dash lines belong to the intermediate-activity category ($50 >$ read count $\geq 10$), below the purple dash line belong to low-activity category (read count $< 10$). Reads from different *msh1* individuals are merged. Read counts are normalized to 10000 total mt genome-mapped reads per sample. Signals from high- and intermediate-activity repeats are labeled.

**Figure S6** Parental forms and recombination products can be detected according to paths
Graph representation of the effect of crossover recombination in a hypothetical circle containing only one pair of direct or inverted repeats. Paths of parental forms (1-1, 2-2) and recombination products (1-2, 2-1) are indicated.

**Figure S7** Two-rearrangement reads involving large repeats are increased in *msh1* mutants
(a-d) Plots show global patterns of the junctions and relative read counts of mitochondrial two-rearrangement reads involving large repeats in (a, c) wild-type and (b, d) *msh1* mutant. Signals from previously named repeats are labeled. Reads from different *msh1* individuals are merged. There are 1668 two-rearrangement reads out of 24162 total reads in all *msh1* samples. Colors represent categories of AO length. The positions of dots are similarly defined as in Figure 2a, but with a constant size. The thickness of lines linking two dots denotes read counts normalized to 10000 total mt genome-mapped reads per sample (line color is the same as the larger dot).

**Figure S8** Continuous distribution of SNVs in non-identical repeats CC-1/2
Breakpoints in SNVs in one-rearrangement HiFi reads indicate possible crossover sites in recombination events between repeat CC-1 and CC-2 repeats in the mt genome of *msh1* mutants. Numbers in brackets denote the sum of read counts in all samples without normalization. The shaded region denotes the longest identical fragment (LIF) in CC repeats. Numbers in parentheses denote the sum of read counts in all samples without normalization. The top two lines are to scale with vertical bars indicating bases of the same colors as the letters. The lines below are not drawn to scale to avoid crowding. Vertical bars in panels a and c denote A (green), T (red), G (brown) and C (blue) for SNVs.

**Figure S9** Mapping of zero-rearrangement long reads identifies variants within repeats

The schematic shows mapping of zero-rearrangement reads to two non-identical repeats with one SNV. Both long and short reads with reference SNVs/indels in the repeated sequence (reference reads) are correctly mapped. Long reads with non-allelic SNVs/indels (NA reads) are correctly mapped based on the long flanking sequences, which indicate potential gene conversion events. Short NA reads fail to identify non-allelic SNVs/indels because of incorrect mapping to the other repeat homolog.

**Figure S10** Imperfect repeats with relatively higher ratios of exchange of SNVs and indels in *msh1* mutants
Square-shaped stacked bar plots are proportions of non-allelic SNVs/indels at predicted sites in repeat group A, D, G, I, M, MMJS, N, R, T and W, which show relatively high accumulation ratios in samples of *msh1* mutants compared to wild-type (W3-5-2, this study; XJTU, Wang et al., 2021; CEN, Naish et al., 2021). The relative heights in each square-shaped box indicate the ratios of given bases for A (green), T (red), G (brown), C (blue), and I (gray) for reference (gray) or non-allelic (purple) sequences at indel sites. Reference bases are indicated below. Results were calculated from zero-rearrangement reads. SNVs and indels are arranged from small to large coordinates in the reference. Coordinates of the first and last SNVs/INDs (black triangles) are shown.

**Figure S11** Coverage of regions containing repeat group H in wild-type and *msh1* individuals
(a-f) Screenshots of (Integrative Genomics Viewer) IGV indicate coverages (a, c, e) all reads and (b, d, f) zero-rearrangement reads of wild-type (green) and *msh1* (brown) individuals in mt genome region (a, b) 31500-32500, (c, d) 269500-270500, and (e, f) 332000-333000 containing repeat H-1, H-2, and H-3 respectively.

**Figure S12** Coverage of regions containing repeat group MMJS in wild-type and *msh1* individuals
(a-d) Screenshots of (Integrative Genomics Viewer) IGV indicate coverages (a, c) all reads and (b, d) zero-rearrangement reads of wild-type (green) and *msh1* (brown) individuals in mt genome region at positions (a, b) 134300-135300, and (c, d) 257300-258300 containing repeat MMJS-1 and MMJS-2 respectively.

**Figure S13** metaFlye assemblies of mt genomes in other accessions
(a, b) Plots show assembly graphs of mt genome-mappped reads of seven *Arabidopsis thaliana* accessions from published PacBio CLR datasets (Jiao and Schneeberger, 2020) with metaFlye for

parameters (a) --genome-size 370K and (b) 2M. In A and B, before_rr denotes graphs before repeat resolution; after_rr denotes graphs after repeat_resolution.

**Figure S14** The metaFlye assemblies of mt genomes are more complex in *msh1* mutants than wild-type

(a, b) Plots show assembly graphs of mt genome-mappped reads of wild-type and *msh1* mutants with metaFlye for parameters (a) --genome-size 370K and (b) 2M. In A and B, before_rr denotes graphs before repeat resolution; after_rr denotes graphs after repeat_resolution. Reads from different *msh1* mutant families are merged. (c, d) Plots show assembly graphs of mt genome-mappped reads of wild-type datasets Col-XJTU (Wang et al., 2021) and Col-CEN (Naish et al., 2021) with metaFlye for parameters (c) --genome-size 370K and (d) 2M. In panels a to d, before_rr denotes graphs before repeat resolution; after_rr denotes graphs after repeat_resolution.

**Figure S15** Synteny analysis of mt genomes of other *Arabidopsis thaliana* accessions

Circos plots show synteny of mitochondrial pseudo-master circles between other *Arabidopsis thaliana* accessions and the Col-0 reference. Direct (grey) and inverted (yellow) synteny are based on blastn alignments. Large repeats in direct (brown) and inverted (red) orientation are shown. Black solid arrowheads denote sequences that are absent in Col-0. Black hollow arrowheads denote sequences that are present in Col-0 but not in the other accession. The repeats that mediate rearrangements and the sizes of large indels are labeled.

**Figure S16** Normalized per base read coverage across mt genome

Plots show normalized per-base read coverage across the entire mt genome of each individual and the published datasets. Annotations are based on the GenBank file NC_037304.1. Purple and orange lines denote normalized per-base read coverage of M3-17 and M3-43 mutant individuals. Green lines denote normalized per-base read coverage of W3-5-2(this study), and the published datasets Col-XJTU (Wang et al., 2021) and Col-CEN (Naish et al., 2021). Red horizontal lines denote 100% coverage.

**Figure S17** Coverages around *atp6-1*, *atp6-2*, and *atp9* in *msh1* mutants and Arabidopsis accessions

(a-c) Plots are normalized per-base read coverage of mt genome regions: positions (a) 49201-51200, (b) 263901-265900, and (c) 270501-271500. Purple and orange lines denote normalized per-base read coverage of M3-17 and M3-43 mutant individuals. Green lines denote normalized per-base read coverage of W3-5-2 (this study), and the published datasets Col-XJTU (Wang et al., 2021) and Col-CEN (Naish et al., 2021). (d-f) Plots are normalized per-base read coverage of mt genome regions: positions (d) 49201-51200, (e) 263901-265900, and (f) 270501-271500. Green lines denote normalized per-base read coverage from the published Col-0 CLR dataset. Purple lines denote normalized per-base read coverage from the published Eri-1 and C24 datasets. Orange lines denote normalized per-base read coverage from the published Cvi-0, An-1, L*er*, Kyoto and Sha datasets. In panels a to f, blue arrows denote coding regions of *atp6-1*, *atp6-2* and *atp9*; purple arrows denote tRNA and other protein-coding genes; orange arrows denote different mitochondrial repeats; annotations are based on the GenBank accession NC_037304.1; red horizontal lines in c to e denote 100% coverage. The jumps in coverage lines were sometimes at repeat boundaries, suggesting results of repeat-mediated rearrangements.

**Figure S18** The proportions of structural variants in Arabidopsis pt genomes

(a) Plots show proportions of different types of pt genome rearrangements detected by HiFi reads in wild-type and *msh1* mutants. The percentages of rearrangement categories are shown in parentheses. The total circles represent all reads, and the minor circles represent the last 2% of total reads to show low-frequency categories. (b) The distribution of Alignment overlap length (AO length) in plastid one-rearrangement events detected by HiFi reads. Counts from read groups with equal AO lengths are summed. Reads from different *msh1* individuals are merged. Read counts are normalized to 100000 total pt genome-mapped reads per sample.

**Figure S19** Length variants at simple sequence repeats in the pt genomes

Heatmap shows the ratios of non-reference lengths at 85 simple sequence repeats of individual samples minus the mean ratios of three wild-type samples. The proportions of length variants at most positions were much higher in *msh1* mutants compared with wild-type, which supported the previously proposed role of *MSH1* in maintaining low mutation rates.
