## Supplemental Figures for "Long-read sequencing characterizes mitochondrial and plastid genome variants in Arabidopsis *msh1* mutants"

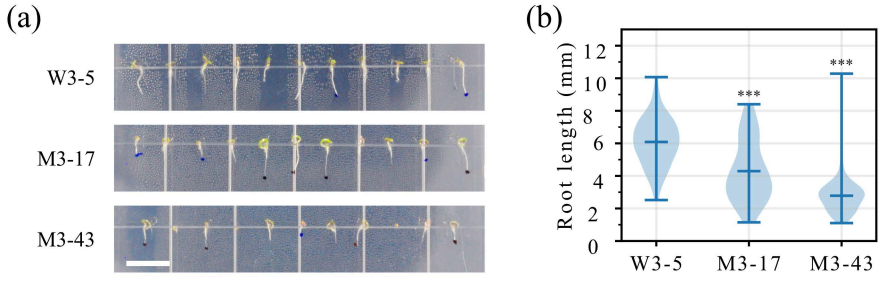
(a)
W3-5
M3-17
M3-43
(b)
Root length (mm)
12
10
8
6
4
2
0
***
***
W3-5
M3-17
M3-43

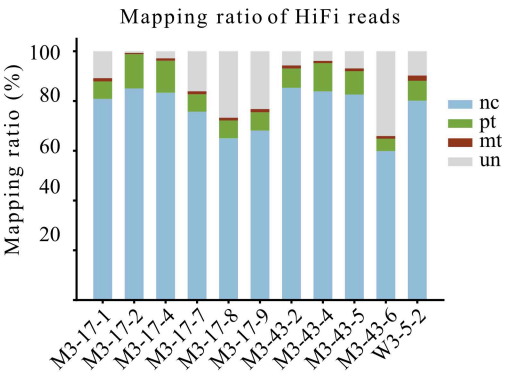
Mapping ratio of HiFi reads
Mapping ratio (%)
100
80
60
40
20
nc
pt
mt
un
M3-17-1
M3-17-2
M3-17-4
M3-17-7
M3-17-8
M3-17-9
M3-43-2
M3-43-4
M3-43-5
M3-43-6
W3-5-2

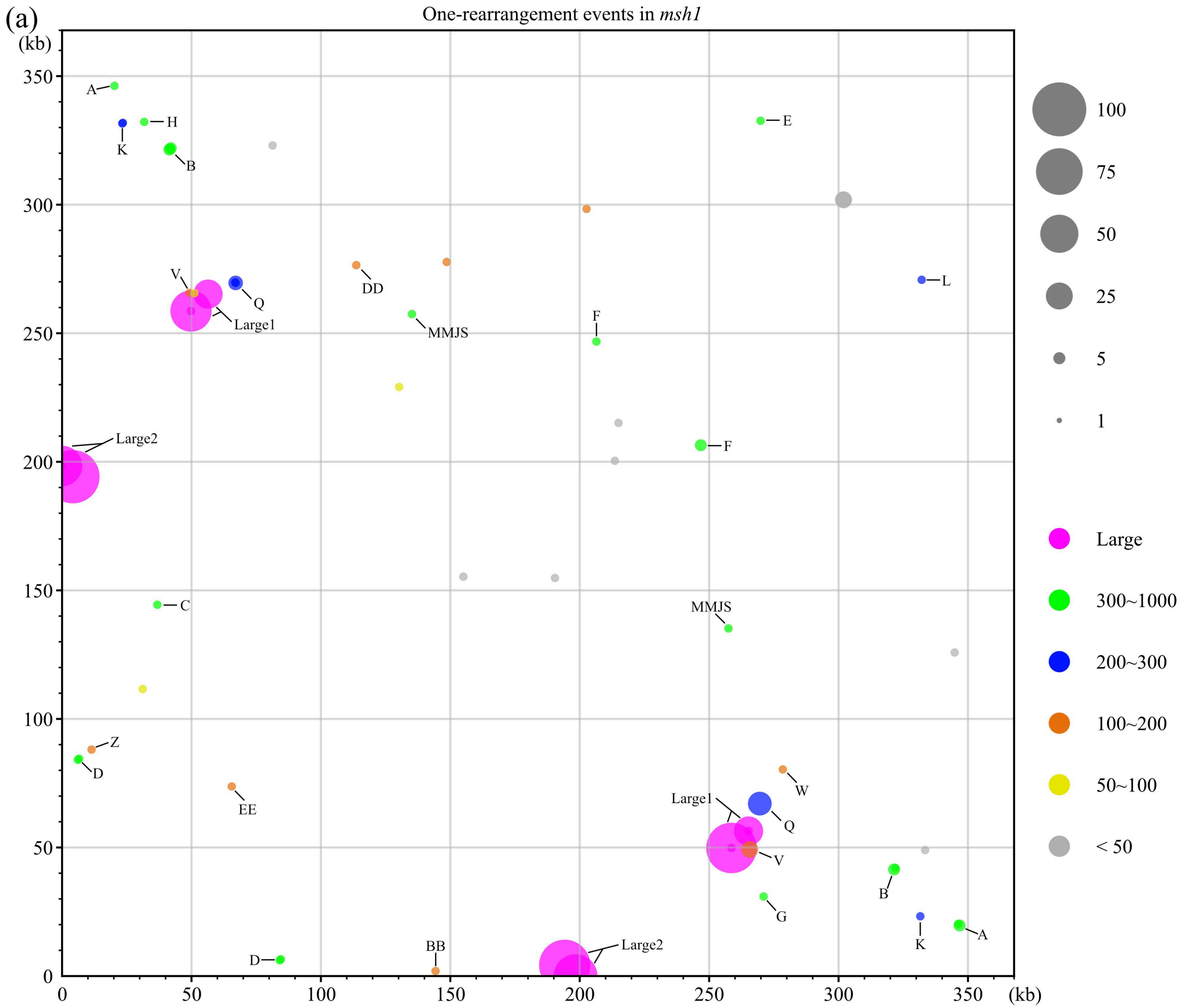

(a)
One-rearrangement events in msh1
(kb)
350
300
250
200
150
100
50
0
50
100
150
200
250
300
350
(kb)
A
H
K
B
E
V
Q
Large1
DD
MMJS
F
L
Large2
F
C
MMJS
Z
D
EE
W
Large1
Q
V
B
G
K
A
D
BB
Large2
100
75
50
25
5
1
Large
300~1000
200~300
100~200
50~100
< 50

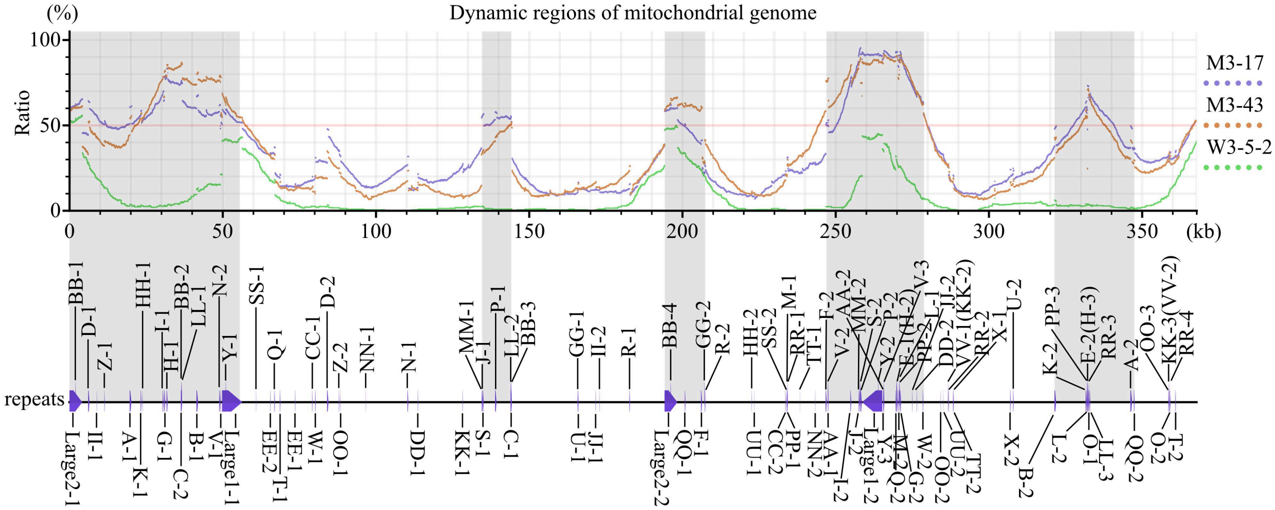

Dynamic regions of mitochondrial genome
(%)
Ratio
100
50
0
0
50
100
150
200
250
300
350
(kb)
M3-17
M3-43
W3-5-2
repeats

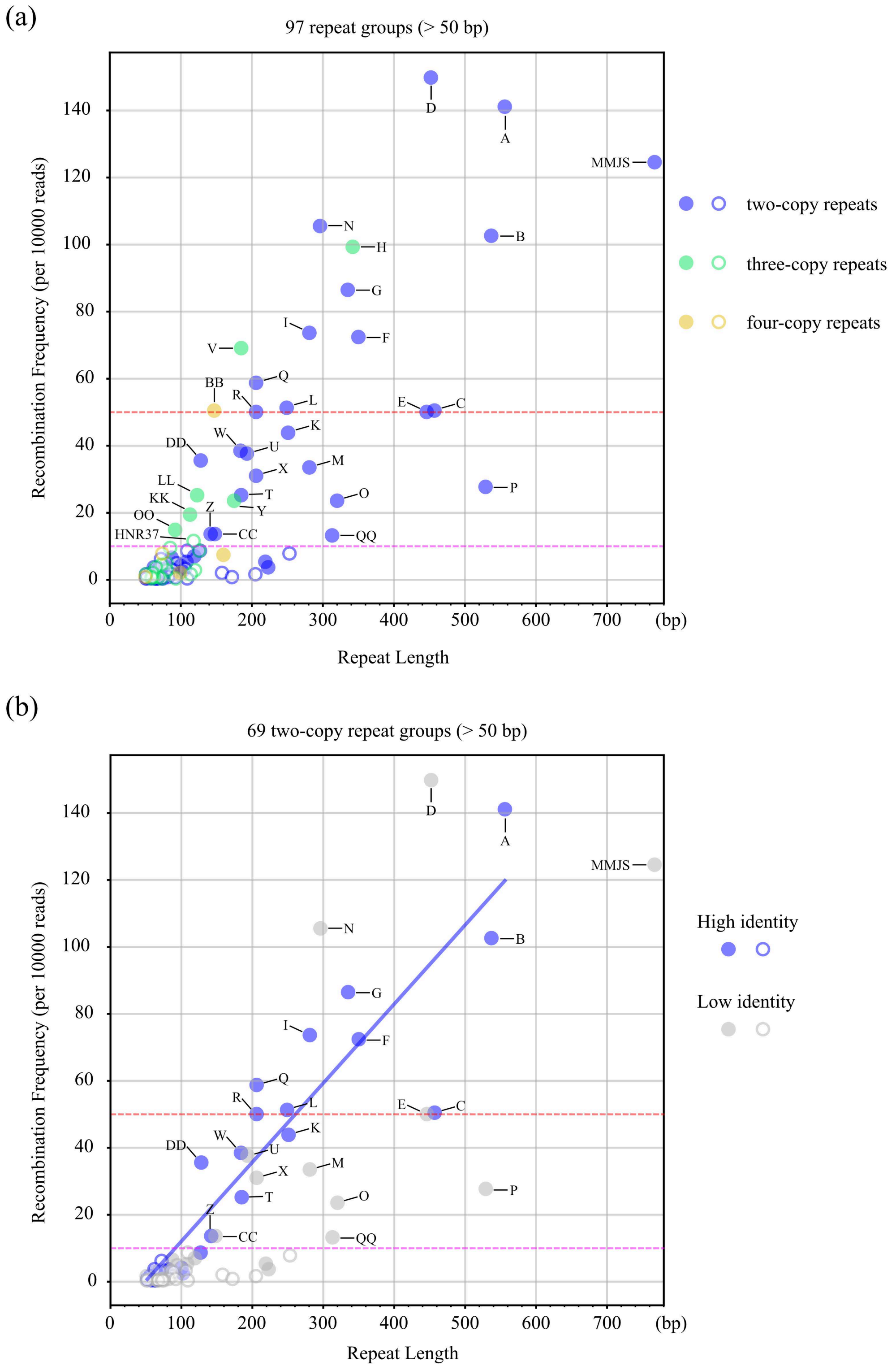
(a)
97 repeat groups (> 50 bp)
Recombination Frequency (per 10000 reads)
Repeat Length
(bp)
two-copy repeats
three-copy repeats
four-copy repeats
D
A
MMJS
N
H
B
G
I
F
V
Q
BB
R
L
E
C
K
DD
W
U
X
M
LL
T
O
P
KK
Y
Z
OO
HNR37
CC
QQ
(b)
69 two-copy repeat groups (> 50 bp)
Recombination Frequency (per 10000 reads)
Repeat Length
(bp)
High identity
Low identity
D
A
MMJS
N
B
G
I
F
Q
R
L
E
C
K
DD
W
U
X
M
T
O
P
Z
CC
QQ

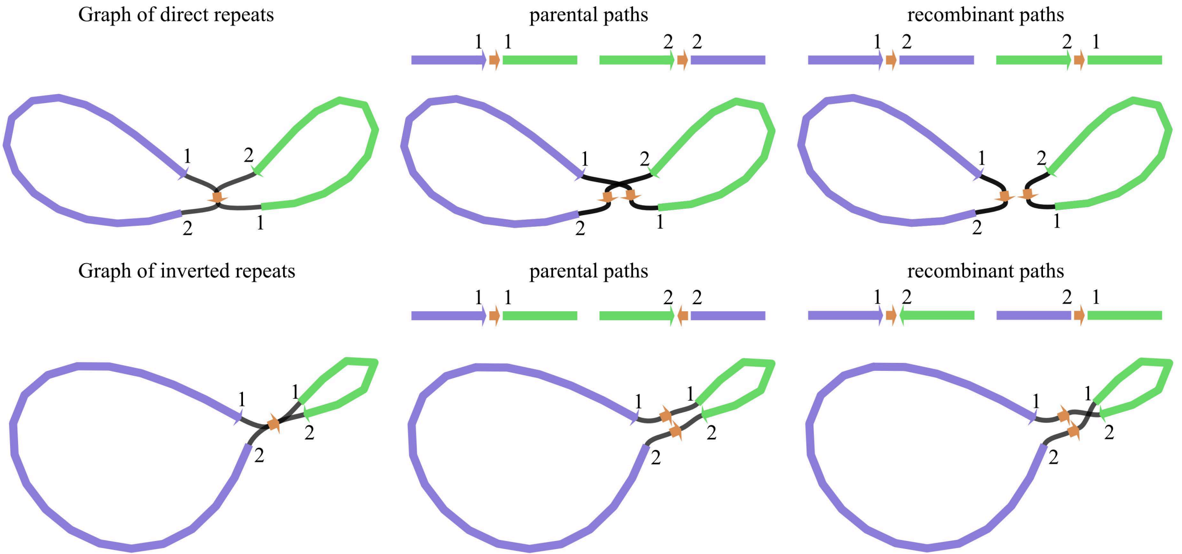

Graph of direct repeats
parental paths
recombinant paths
Graph of inverted repeats
parental paths
recombinant paths
1
2

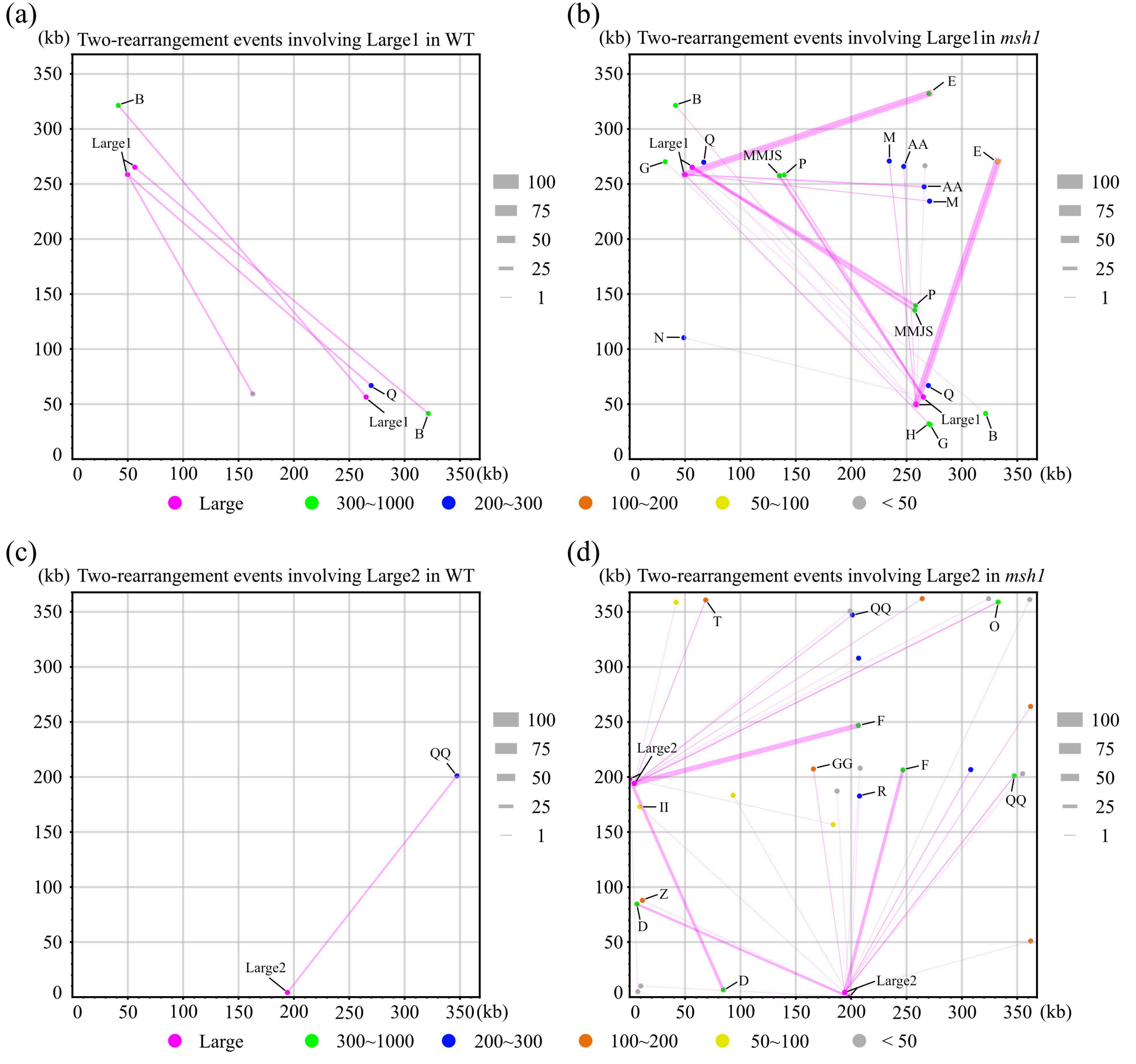
(a)
(kb) Two-rearrangement events involving Large1 in WT
(b)
(kb) Two-rearrangement events involving Large1in msh1
(c)
(kb) Two-rearrangement events involving Large2 in WT
(d)
(kb) Two-rearrangement events involving Large2 in msh1
Large
300~1000
200~300
100~200
50~100
< 50
100
75
50
25
1

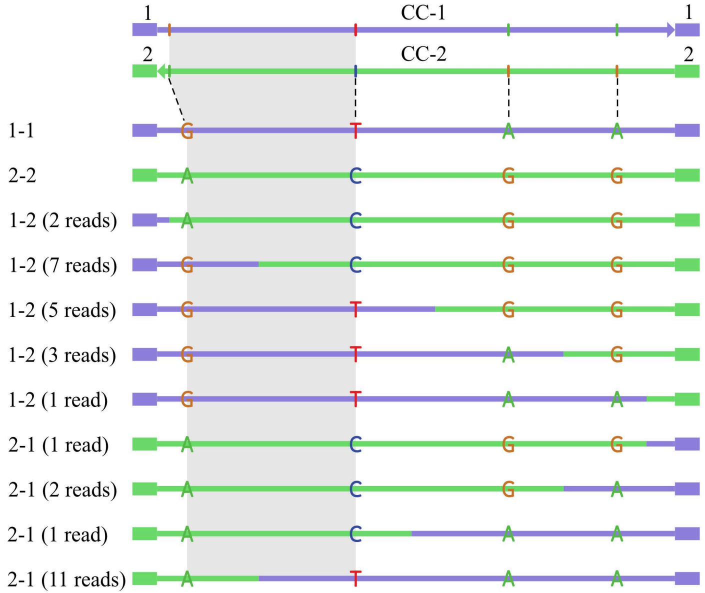

1
CC-1
1
2
CC-2
2
1-1
G T A A
2-2
A C G G
1-2 (2 reads)
A C G G
1-2 (7 reads)
G C G G
1-2 (5 reads)
G T G G
1-2 (3 reads)
G T A G
1-2 (1 read)
G T A A
2-1 (1 read)
A C G G
2-1 (2 reads)
A C G A
2-1 (1 read)
A C A A
2-1 (11 reads)
A T A A

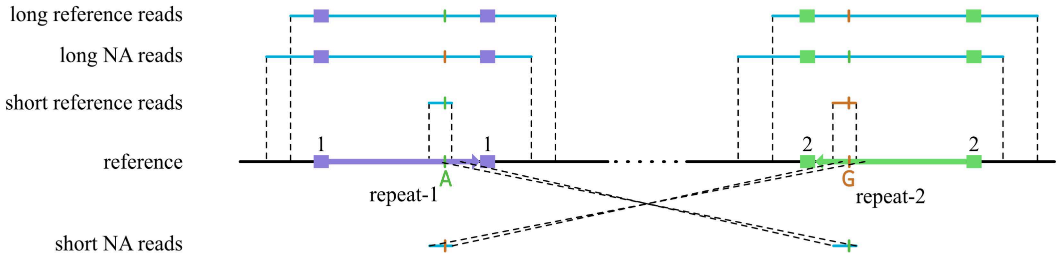
long reference reads
long NA reads
short reference reads
reference
short NA reads
1
1
2
2
A
G
repeat-1
repeat-2

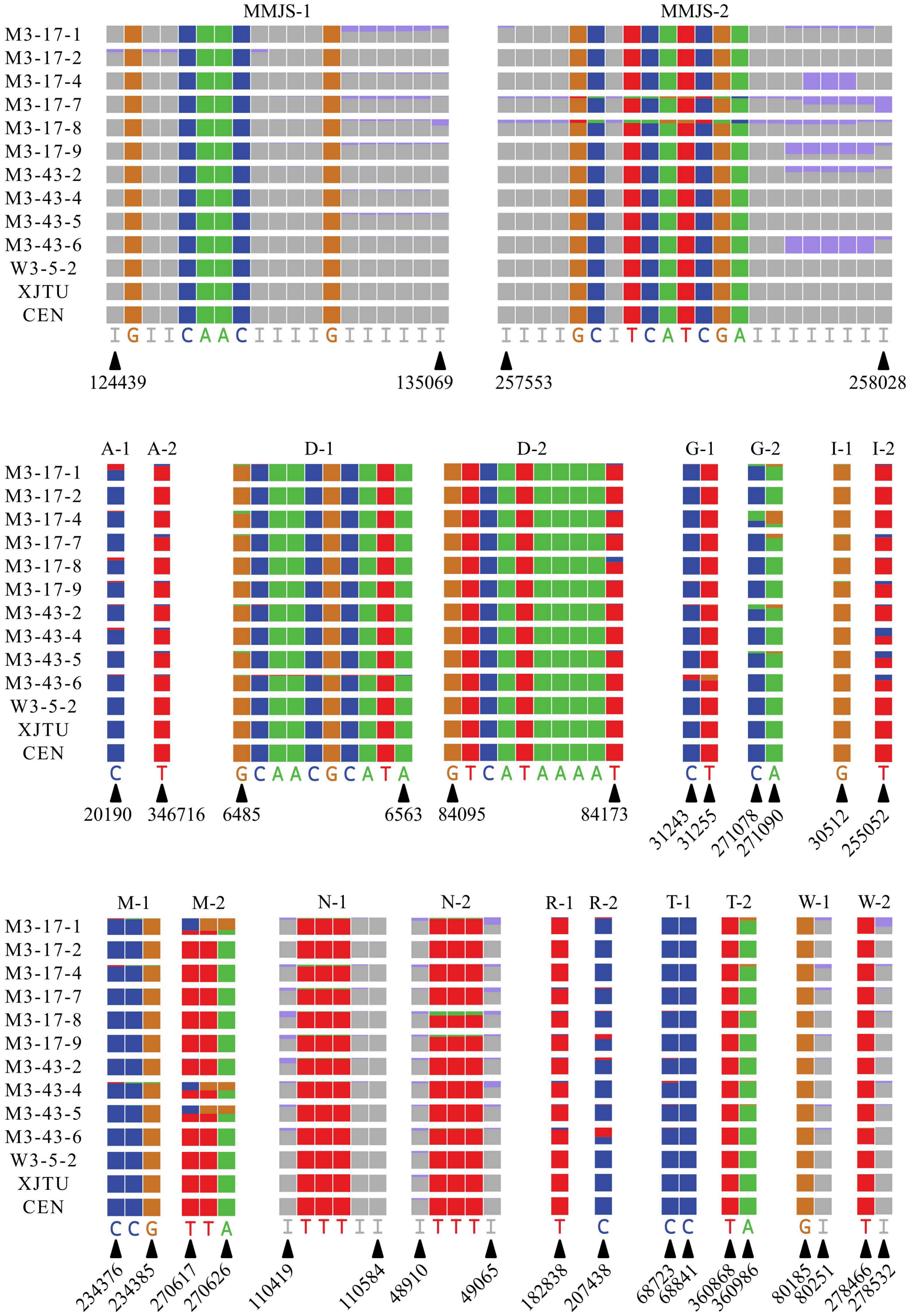

MMJS-1
MMJS-2
M3-17-1
M3-17-2
M3-17-4
M3-17-7
M3-17-8
M3-17-9
M3-43-2
M3-43-4
M3-43-5
M3-43-6
W3-5-2
XJTU
CEN
I G I I C A A C I I I I G I I I I I I
I I I I G C I T C A T C G A I I I I I I I I
124439
135069
257553
258028
A-1
A-2
D-1
D-2
G-1
G-2
I-1
I-2
C
T
G C A A C G C A T A
G T C A T A A A A T
C T
C A
G
T
20190
346716
6485
6563
84095
84173
31243
31255
271078
271090
30512
255052
M-1
M-2
N-1
N-2
R-1
R-2
T-1
T-2
W-1
W-2
C C G
T T A
I T T T I I
I T T T I
T
C
C C
T A
G I
T I
234376
234385
270617
270626
110419
110584
48910
49065
182838
207438
68723
68841
360868
360986
80185
80251
278466
278532

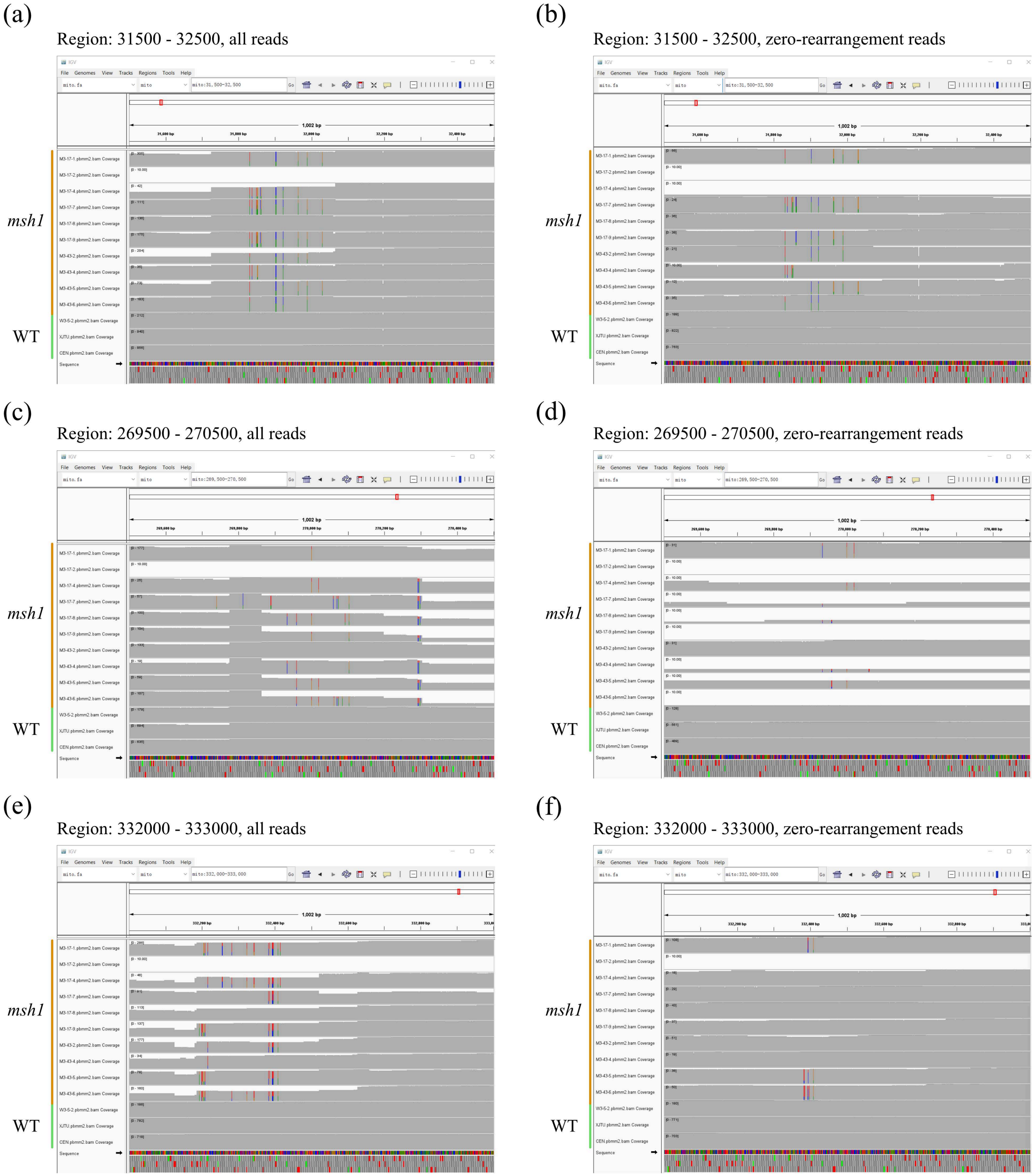
(a)
Region: 31500 - 32500, all reads
(b)
Region: 31500 - 32500, zero-rearrangement reads
(c)
Region: 269500 - 270500, all reads
(d)
Region: 269500 - 270500, zero-rearrangement reads
(e)
Region: 332000 - 333000, all reads
(f)
Region: 332000 - 333000, zero-rearrangement reads
msh1
WT

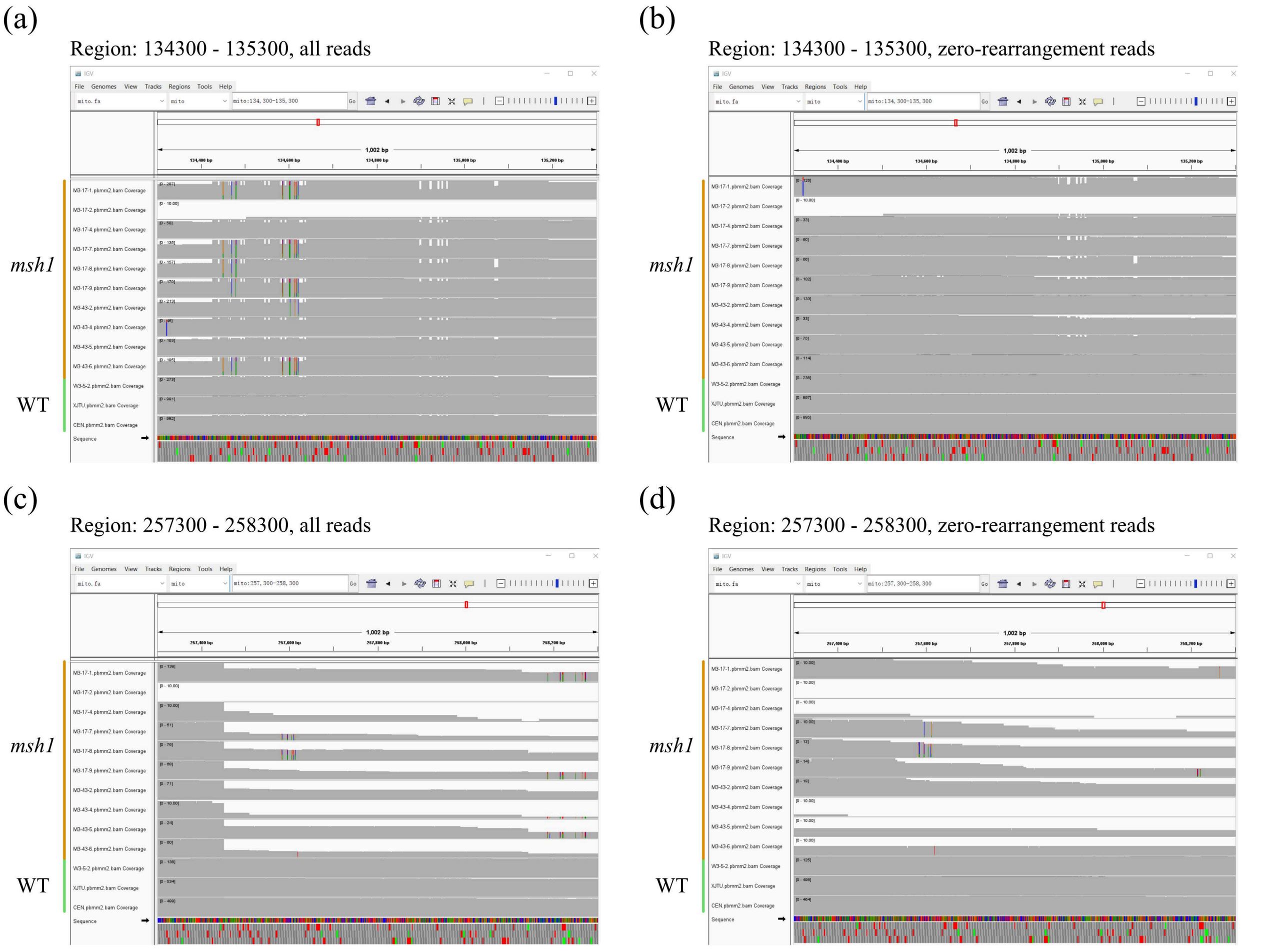
(a)
Region: 134300 - 135300, all reads
(b)
Region: 134300 - 135300, zero-rearrangement reads
(c)
Region: 257300 - 258300, all reads
(d)
Region: 257300 - 258300, zero-rearrangement reads
msh1
WT

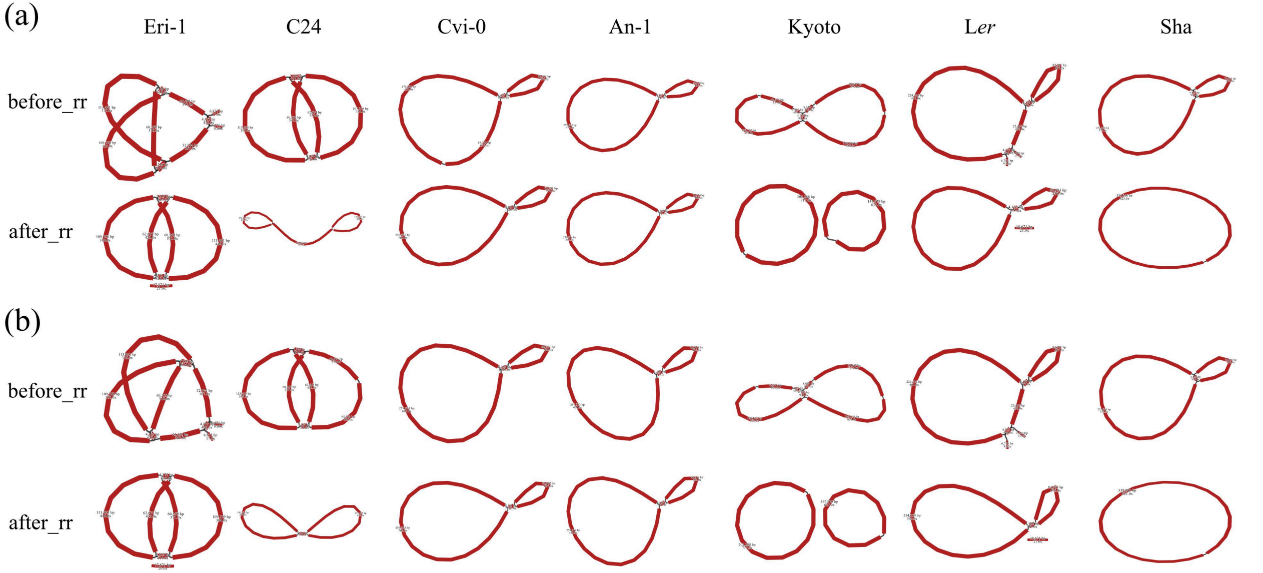
(a)
Eri-1
C24
Cvi-0
An-1
Kyoto
Ler
Sha
before_rr
after_rr
(b)
before_rr
after_rr

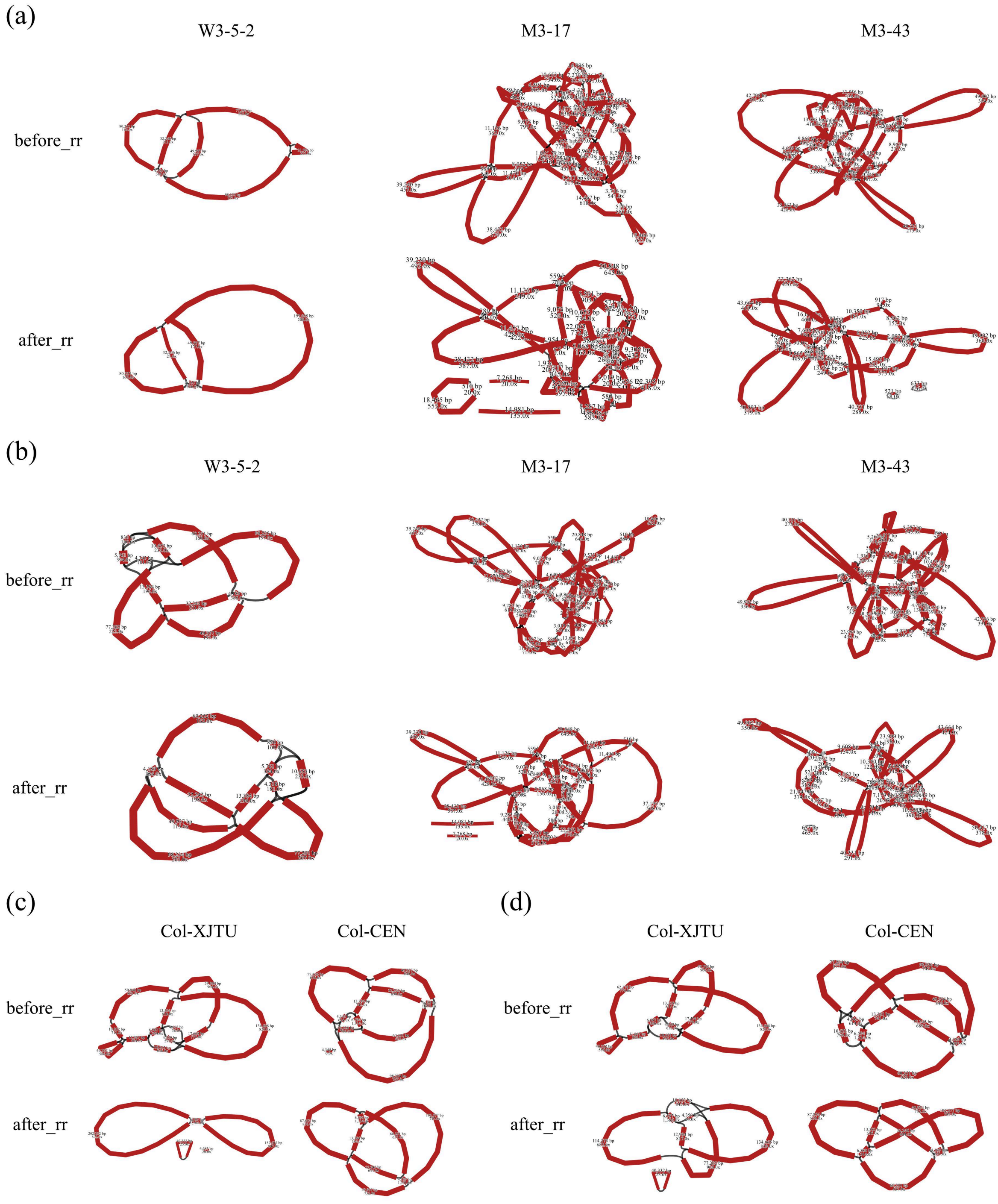

(a)
W3-5-2
M3-17
M3-43
before_rr
after_rr
(b)
W3-5-2
M3-17
M3-43
before_rr
after_rr
(c)
Col-XJTU
Col-CEN
before_rr
after_rr
(d)
Col-XJTU
Col-CEN
before_rr
after_rr

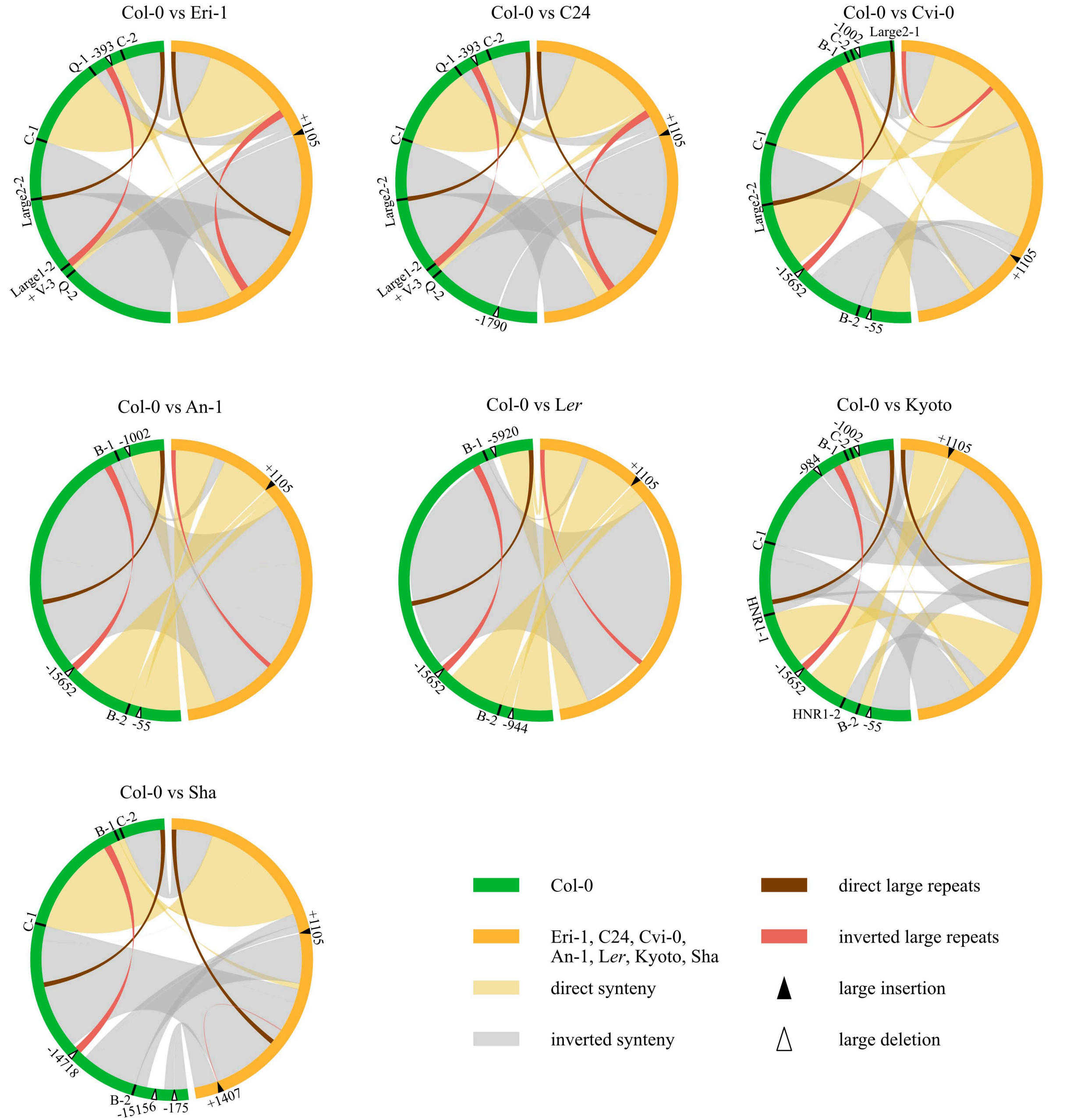

Col-0 vs Eri-1
Q-1
-393
C-2
C-1
Large2-2
Large1-2
+ V-3
Q-2
+1105
Col-0 vs C24
Q-1
-393
C-2
C-1
Large2-2
Large1-2
+ V-3
Q-2
-1790
+1105
Col-0 vs Cvi-0
-1002
C-2
B-1
Large2-1
C-1
Large2-2
-15652
B-2
-55
+1105
Col-0 vs An-1
B-1
-1002
+1105
-15652
B-2
-55
Col-0 vs Ler
B-1
-5920
+1105
-15652
B-2
-944
Col-0 vs Kyoto
-1002
C-2
B-1
-984
+1105
C-1
HNR1-1
-15652
HNR1-2
B-2
-55
Col-0 vs Sha
B-1
C-2
C-1
+1105
-14718
B-2
-15156
-175
+1407
Col-0
Eri-1, C24, Cvi-0,
An-1, Ler, Kyoto, Sha
direct synteny
inverted synteny
direct large repeats
inverted large repeats
large insertion
large deletion

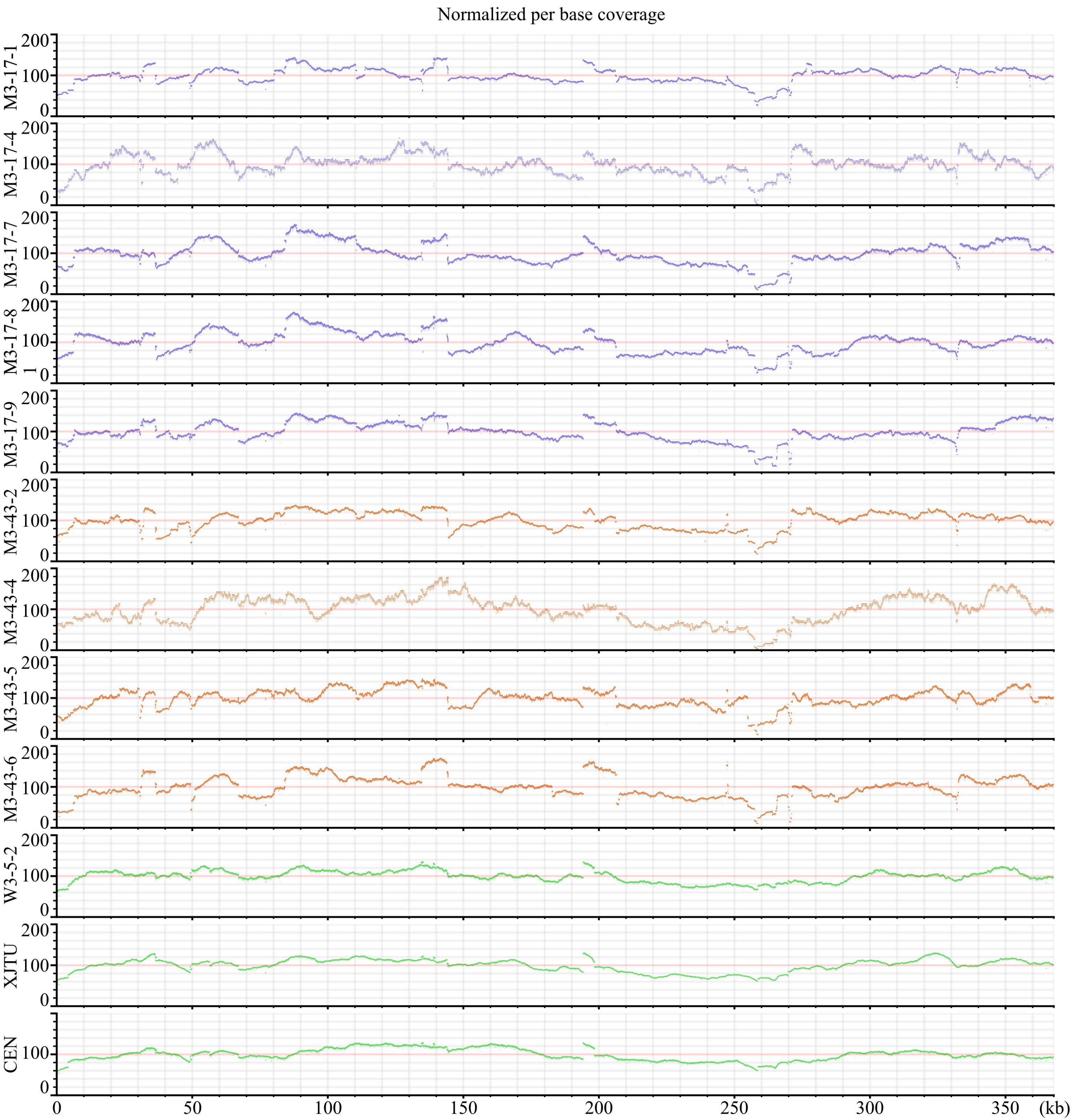

Normalized per base coverage
M3-17-1
M3-17-4
M3-17-7
M3-17-8
M3-17-9
M3-43-2
M3-43-4
M3-43-5
M3-43-6
W3-5-2
XJTU
CEN
0
50
100
150
200
250
300
350
(kb)

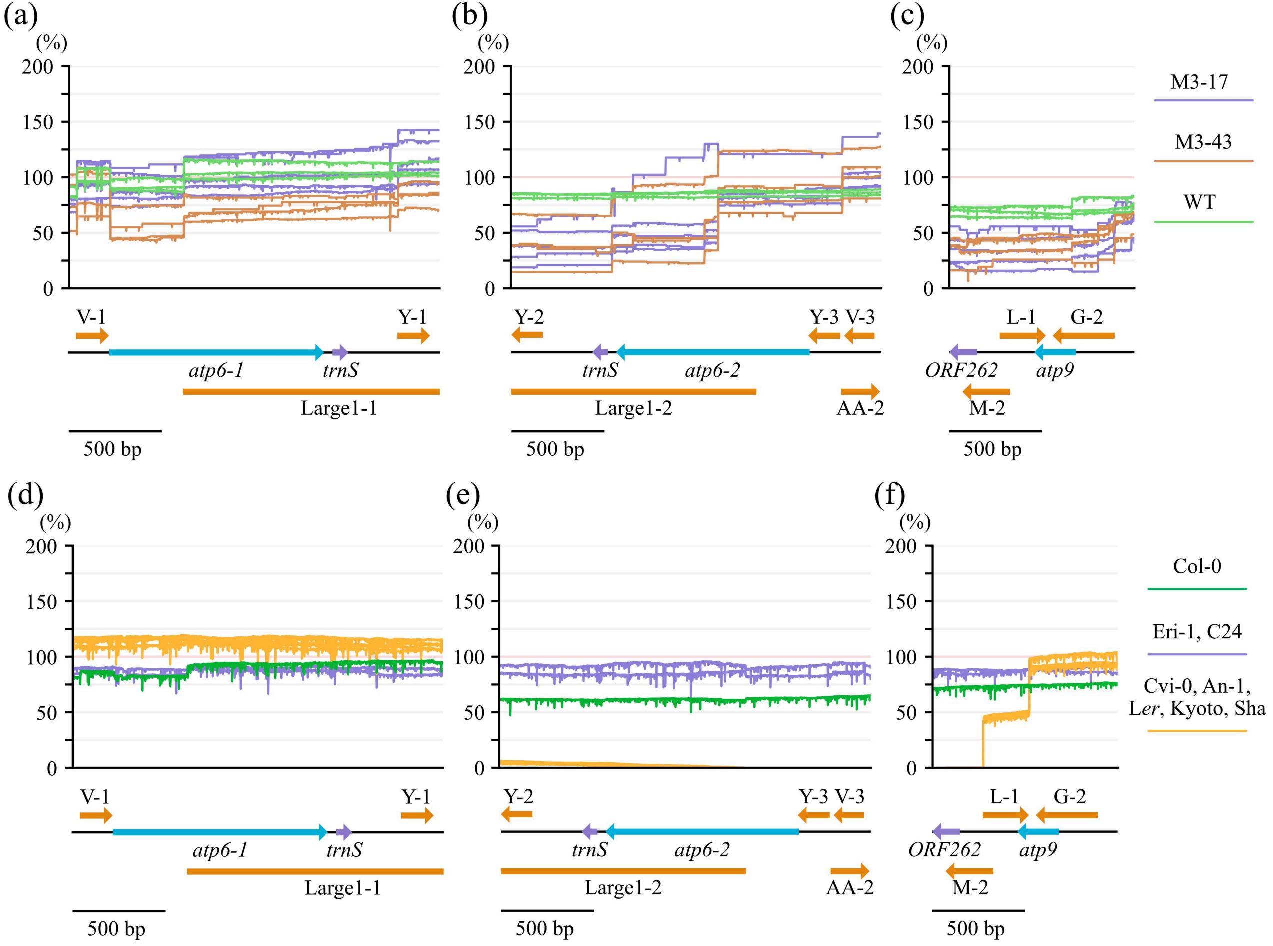

(a)
(b)
(c)
(%)
200
150
100
50
0
V-1
Y-1
atp6-1
trnS
Large1-1
500 bp
Y-2
Y-3
V-3
trnS
atp6-2
Large1-2
AA-2
L-1
G-2
ORF262
atp9
M-2
M3-17
M3-43
WT
(d)
(e)
(f)
Col-0
Eri-1, C24
Cvi-0, An-1,
Ler, Kyoto, Sha

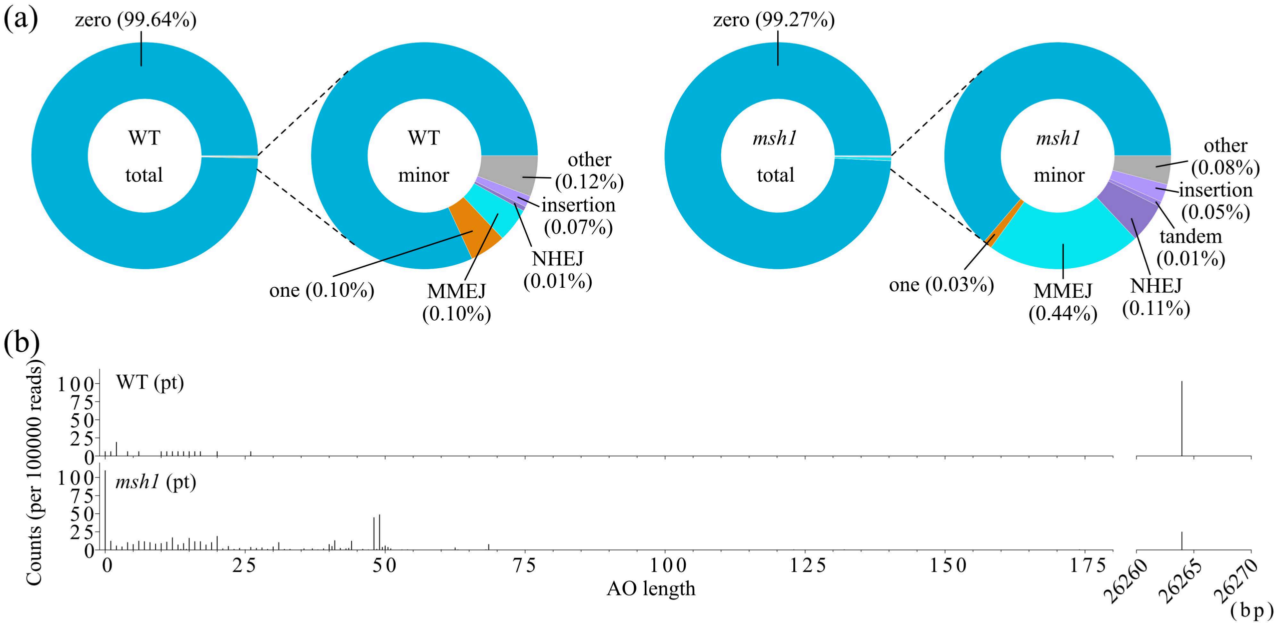

(a)
zero (99.64%)
WT
total
WT
minor
other
(0.12%)
insertion
(0.07%)
NHEJ
(0.01%)
MMEJ
(0.10%)
one (0.10%)
zero (99.27%)
msh1
total
msh1
minor
other
(0.08%)
insertion
(0.05%)
tandem
(0.01%)
NHEJ
(0.11%)
MMEJ
(0.44%)
one (0.03%)
(b)
WT (pt)
msh1 (pt)
Counts (per 100000 reads)
0
25
50
75
100
0
25
50
75
100
125
150
175
26260
26265
26270
AO length
(bp)

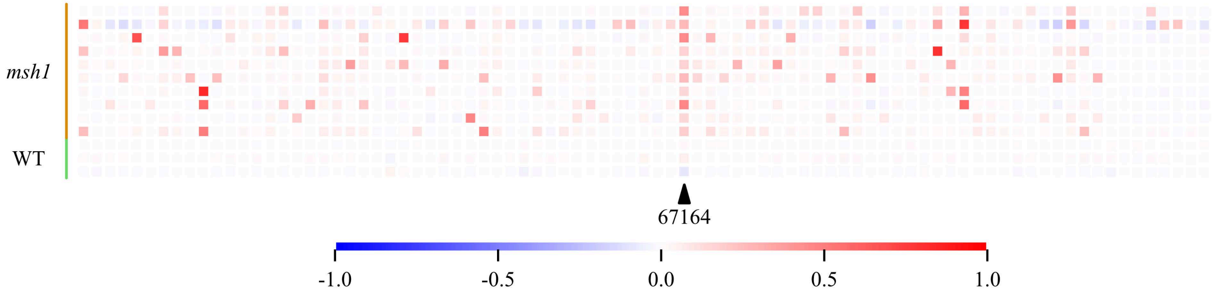
msh1
WT
67164
-1.0
-0.5
0.0
0.5
1.0
